# miR-378a Controls Cardiomyocyte Metabolism and Angiogenic Signaling

**DOI:** 10.64898/2026.06.23.733812

**Authors:** Jacek Stępniewski, Alicja Martyniak, Izabella Więckowska, Tomasz Gaczorek, Gabriela Machaj, Ewelina Pośpiech, Luisa Schmidt, Theresa Bock, Mateusz Tomczyk, Izabela Kraszewska, Katarzyna Sarad, Joanna Korytowska, Katarzyna Polak, Natalia Limberger, Olga Barczyk-Woźnicka, Elżbieta Pyza, Marcus Krüger, Guillem Ylla, Mauro Giacca, Józef Dulak, Urszula Florczyk-Soluch

## Abstract

**Aims:** While the muscle-enriched microRNA-378a (miR-378a) has been implicated in cardiac hypertrophy and stress responses, its role in maintaining cardiomyocyte metabolic homeostasis, mitochondrial function, and angiogenic paracrine signaling under physiological and post-injury conditions remains unclear. This study addresses these gaps by examining the molecular and functional consequences of miR-378a deficiency in murine heart and human cardiomyocytes.

**Methods and Results:** Cardiac structure and function were analyzed in miR-378a-deficient (miR-378a−/−) and wild-type (miR-378a+/+) mice at 12 weeks and 17 months of age, revealing that miR-378a loss promoted myocardial fibrosis, altered IGF1R–AKT signaling, and impaired cardiac performance, with age-dependent effects.

Integrated transcriptomic and proteomic analyses in miR-378a-/- and control mice, as well as in human iPSC-derived cardiomyocytes (hiPSC-CM) of both genotypes, revealed deregulated pathways related to translation, metabolism, and cardiomyopathy-associated signaling. In hiPSC-CM, miR-378a knockout (KO) impaired mitochondrial respiration, disrupted mitochondrial morphology, and reduced mitochondrial DNA content, accompanied by altered mitophagy and biogenesis. KO cells also showed increased glucose uptake but reduced glycogen storage, accompanied by changes in key metabolic regulators, and displayed diminished angiogenic potential. Finally, hiPSC-CM overexpressing miR-378a were delivered in a mouse model of acute myocardial infarction, but overexpression did not further enhance their therapeutic effect.

**Conclusions:** This study broadens our understanding of miR-378a’s physiological role in murine hearts and human cardiomyocytes, demonstrating its impact on contractility, mitochondrial integrity, glucose metabolism, and angiogenic paracrine signaling. However, overexpression of miR-378a in hiPSC-CM offers limited additional benefit in cell therapy for acute myocardial infarction.

## 1. Introduction

Within the cardiovascular system, microRNAs (miRNAs) are abundantly expressed and have been shown to play crucial roles in, among others, cell differentiation, proliferation, apoptosis, and angiogenesis (1). Among them, muscle-enriched miR-378a has attracted particular interest due to its roles in energy metabolism and its potential impact on vascular processes (2–4). Notably, miR-378a is predominantly expressed in cardiomyocytes, increasing over time in both animal models (5) and during long-term culture of human embryonic stem cell-derived cardiomyocytes (hESC-CM) (6).

Most research on cardiac miR-378a focuses on pathological conditions, such as chronic hypertension, hypertrophy and hypoxia. Ganesan et al. (7) reported that miR-378a is downregulated in murine models of chronic pressure overload and severe heart failure, mirroring reduced levels in human hearts with dilated cardiomyopathy. In neonatal rat cardiomyocytes, miR-378a targets IGF1R and downstream MAPK pathway components, controlling hypertrophic growth (7). We extended these findings to human induced pluripotent stem cell-derived cardiomyocytes (hiPSC-CM) (8), showing that miR-378a-deficient cells exhibited significantly larger size in comparison to the isogenic control group and exhibited higher levels of pro-hypertrophic factors, including NFATc3, phospho-AKT, phospho-ERK and ERK (8).

Beyond hypertrophy, miR-378a modulates metabolism and confers cytoprotection in cardiomyocytes. In doxorubicin-induced cardiotoxicity, miR-378a overexpression reduced ER stress, lowered a lactate dehydrogenase A (LDHA) levels, preserved mitochondrial membrane potential, and protected neonatal mouse cardiomyocytes from apoptosis (9). In our previous study (10), hyperglycemic miR-378a+/+ mouse hearts and hiPSC-CM exposed to high glucose showed increased miR-378a expression and activation of pro-hypertrophic pathways, suggesting a role in regulating energy homeostasis, glucose metabolism, and mitochondrial function. This aligns with the primary role of miR-378a as a metabolism-modulating factor (2) and its high expression in energetically demanding tissues such as skeletal muscles and myocardium (3).

Despite these insights, miR-378a’s role in maintaining cardiomyocyte function and metabolism under normal physiological conditions, and its contribution after injury, remains incompletely understood. To address this, we performed functional, histological, and proteomic analyses in control and miR-378a-deficient murine hearts, compared the results with transcriptomic and proteomic data from hiPSC-CM, and investigated miR-378a’s role in human cardiomyocyte metabolism. Additionally, we evaluated whether overexpression of miR-378a in hiPSC-CM could enhance their therapeutic potential in a murine model of acute myocardial infarction (MI), using NOD-SCID mice to allow human cell engraftment.

## 2. Materials and Methods

### 2.1. Cell culture

Lines of hiPSC used in the study were generated from peripheral blood mononuclear cells isolated from three healthy donors (two males and one female) as described previously (8), upon obtaining informed consent in accordance with the Declaration of Helsinki and with the approval of the Institutional Review Board and the Bioethical Committee. The cells were cultured on Geltrex^TM^ (ThermoFisher Scientific)-coated 12-well plates in Essential 8 (E8) medium (ThermoFisher Scientific). Medium was refreshed daily, and cells were passaged at ∼70% confluency using 0.5 mM EDTA. Cells were seeded in E8 medium with 10 μM ROCK inhibitor (Y-27632, Abcam) for 24 h.

Primary human aortic endothelial cells (HAEC, obtained from PromoCell at passage 2) were cultured in endothelial cell growth medium (EGM-2, Lonza) composed of EBM-2 Basal Medium and EGM-2 SingleQuots Supplements and used for experiments until passages 5–10.

### 2.2. Animals

The work has been reported in line with the ARRIVE guidelines 2.0. MicroRNA-378-/- mice on a 129SvEv/C57BL/6 background were kindly provided by Eric Olson (Department of Molecular Biology, University of Texas Southwestern Medical Center, Dallas, Texas, USA) (2). To establish a pure C57BL/6 genetic background, 129SvEv/C57BL/6 (miR-378-/-) mice were backcrossed with C57BL/6 mice over ten generations. Counterpart C57BL/6 mice (miR-378+/+) were used as controls. NOD.CB-17-Prkdc acid/Rj mice (NOD-SCID) were purchased from Janvier Labs (Le Genest-Saint-Isle, France). Mice were housed under specific pathogen-free (SPF) conditions in individually ventilated cages, following a 14 h light / 10 h dark cycle, and were provided with standard chow diet and water ad libitum.

All experiments were conducted on male mice. Mice were euthanized by CO₂ inhalation, and hearts were subsequently perfused with heparinized saline (0.5 IU/mL) and 30 mM KCl to induce diastolic arrest.

Sample sizes were determined a priori based on previous experience and statistical considerations, taking into account potential complications or the need for early humane endpoint termination. Animals that reached predefined humane endpoints (e.g., symptoms of wasting) were euthanized and excluded from subsequent analyses.

Animals were randomly allocated to experimental groups, with group assignment adjusted to ensure comparable distributions of body weight and age across groups. No study protocol was preregistered. The study was conducted in accordance with approvals from the relevant ethical committees. All experimental procedures, measurements, and data analyses were conducted blinded to group identity, which was revealed afterward. No unexpected adverse events were observed during the study.

### 2.3. Myocardial infarction and administration of hiPSC-CM

To test whether miR-378a enhances the therapeutic efficacy of transplanted hiPSC-CM, NOD-SCID mice at 6-8 weeks were subjected to myocardial infarction (MI) and cell therapy. Mice were anesthetized with intraperitoneal injection of ketamine (75 mg/kg of body weight) and medetomidine (1 mg/kg of body weight). MI was induced by permanent ligation of the left anterior descending coronary artery as previously described (11). hiPSC1-CM-Luc-empty and hiPSC1-CM-Luc-miR-378a at differentiation days 30–35 were collected from culture plates, counted and immediately injected into the myocardium at four sites in the border of the ischemic area, with 2.5 µL of cells (1 × 10^5^ per site) suspended in saline (total 4 × 10^5^ cells in 10 µL). Atipamezole (5 mg/kg, i.p.) was administered to reverse the effects of medetomidine. Animals received analgesia twice daily for 3 consecutive days after surgery via subcutaneous injection of buprenorphine (0.1 mg/kg).

### 2.4. Transthoracic echocardiography (TTE)

To evaluate cardiac function transthoracic echocardiography (TTE) was performed on miR-378a–/– mice at 10-12 weeks and 7 months of age, along with age-matched wild-type controls (miR-378a+/+) as well as on NOD-SCID mice on day 14, 28 and 42 after MI, as described previously (11). Briefly, anaesthesia was induced using 5% isoflurane (Aerrane, Baxter) in air (v/v) and maintained at 2-2.5%. The mice were immobilized in the supine position on a heating platform. Heart rate and respiration were continuously monitored by ECG electrodes. TTE measurement was performed using high-resolution ultrasound imaging system (Vevo 2100, Visual Sonics) with an MS-400 (18-38 MHz) MicroScan transducer. The heart was first imaged in B-mode (2D) in the parasternal long axis (PSLA), and short axis (SAX) at the level of papillary muscles with the M-mode line placed in the centre of the LV. The integrated software tool (LV-Trace) was used for single-plane SAX analysis. For quantification of cardiac function anterior and posterior walls were traced through 3–4 beats on 2–4 independent tracings.

### 2.5. Bioluminescence imaging

To track the survival and engraftment of transplanted hiPSC-CMs in vivo over time, bioluminescence imaging was performed on NOD-SCID mice using an IVIS Lumina II detector on days 7, 14, 28 and 42 after surgery as described previously (11). Mice were anesthetized with 5% v/v isoflurane/air for induction and 1.5–2% v/v isoflurane/air for maintenance.

### 2.6. Seahorse XF Mito Stress

The energetic potential of hiPSC-CM was tested using the Seahorse XF Mito Stress kit (Agilent). 2 × 10^4^ cells were seeded per well of a plate designed for this test (Agilent). Oligomycin (1.5 μg/mL), FCCP (1 μM), rotenone and antimycin A (1.2 μM) were used to inhibit specific complexes of the mitochondrial respiratory chain. The test was performed according to the manufacturer’s instructions. The results of the analysis were normalized based on the measurement of the concentration of protein isolated from cells by overnight incubation in a lysis buffer (RIPA with the addition of protease inhibitors) using the BCA method.

### 2.7. Mitophagic flux analysis

Mitophagy was induced by a 6-hour stimulation with CCCP (20 μM, Sigma-Aldrich), a compound that uncouples the mitochondrial respiratory chain. Stimulation was continued during staining with a fluorescent probe that selectively labels autophagic vacuoles from Autophagy Assay Kit (Abcam), diluted 1:1000 according to the manufacturer’s instructions. After staining, cells were washed twice with PBS, filtered through a 40 μm strainer, and analysed on BD LSRFortessa flow cytometer (BD Biosciences).

### 2.8. Statistics

All statistical analyses were performed using GraphPad Prism 8™ software. For multiple group comparisons, *two-way ANOVA* or *one-way ANOVA* was applied, followed by Tukey’s post hoc test. Two independent groups were compared using an unpaired two-tailed t-test (with Welch’s correction if variances were unequal). For data not meeting normality assumptions, the nonparametric Mann-Whitney U test was used. *p*-values between 0.05 and 0.1 were considered indicative of a statistical tendency.

All animals allocated to the experimental groups were included in the study and analyses. No inclusion or exclusion criteria were defined prior to the experiment. Data exclusion was limited to outlier detection using Grubbs’ test.

The number of mice, cell lines, and repetitions of in vitro experiments are indicated in the figure legends. The number of animals was limited to the minimum required to obtain statistically significant results. Group sizes varied depending on the type of experiment and the variables involved, such as induction of myocardial infarction or administration of modified versus control cells.

All other methods including cardiac differentiation of hiPSC-CM, AAV production, transcriptomic and proteomic analyses, Western blot, reverse transcription, quantitative PCR, immunofluorescent staining are described in detail in the Supplementary Material.

## 3. Results

### 3.1. miR-378a knockout affects murine heart function

The biological role of miR-378a has been mainly studied in the context of cardiomyocyte stress responses and pathological hypertrophy (7, 9). While our recent study extended these observations to diabetic cardiomyopathy (10), its contribution to cardiac physiology under basal conditions, and how this may vary with age, has not been fully characterized.

Histological analysis of 11-12-week-old mice revealed increased collagen deposition in miR-378a-deficient hearts (Fig. 1A,B) corroborating our previous findings (10) and indicating anti-fibrotic activity of miR-378a. Importantly, TTE performed in 10-13 week old mice (representative recording in Supplementary Fig. 1A) demonstrated significantly decreased left ventricle (LV) ejection fraction (EF, Fig. 1C) and fractional shortening (FS, Fig. 1D) followed by increased end-systolic diameter (Fig. 1E) and volume (Supplementary Fig. 1A) in miR-378a-/- animals in comparison to their control counterparts. At the same time, modest differences (p>0.05) in other cardiac parameters, i.e., end-diastolic diameter (Fig. 1F), end-diastolic volume (Supplementary Fig. 1A) and stroke volume (SV, Supplementary Fig. 1A) were detected. On the other hand, although the level of inflammation and necrosis was slightly higher in miR-378a-/- specimens, the difference did not reach statistical significance (Supplementary Fig. 1B). Further molecular analysis revealed no differences in VEGFA level in heart lysates of both genotypes (Supplementary Fig. 1C) while the level of FGF2 was decreased in miR-378a-deficient samples (Supplementary Fig. 1D). In parallel, immunohistochemical staining demonstrated a slight decrease in the number of αSMA-, CD31- and lectin-positive vessels in miR-378a-/- hearts in comparison to their wild type counterparts (Supplementary Fig. 1E), however this difference did not reach statistical significance.

**Figure 1.**
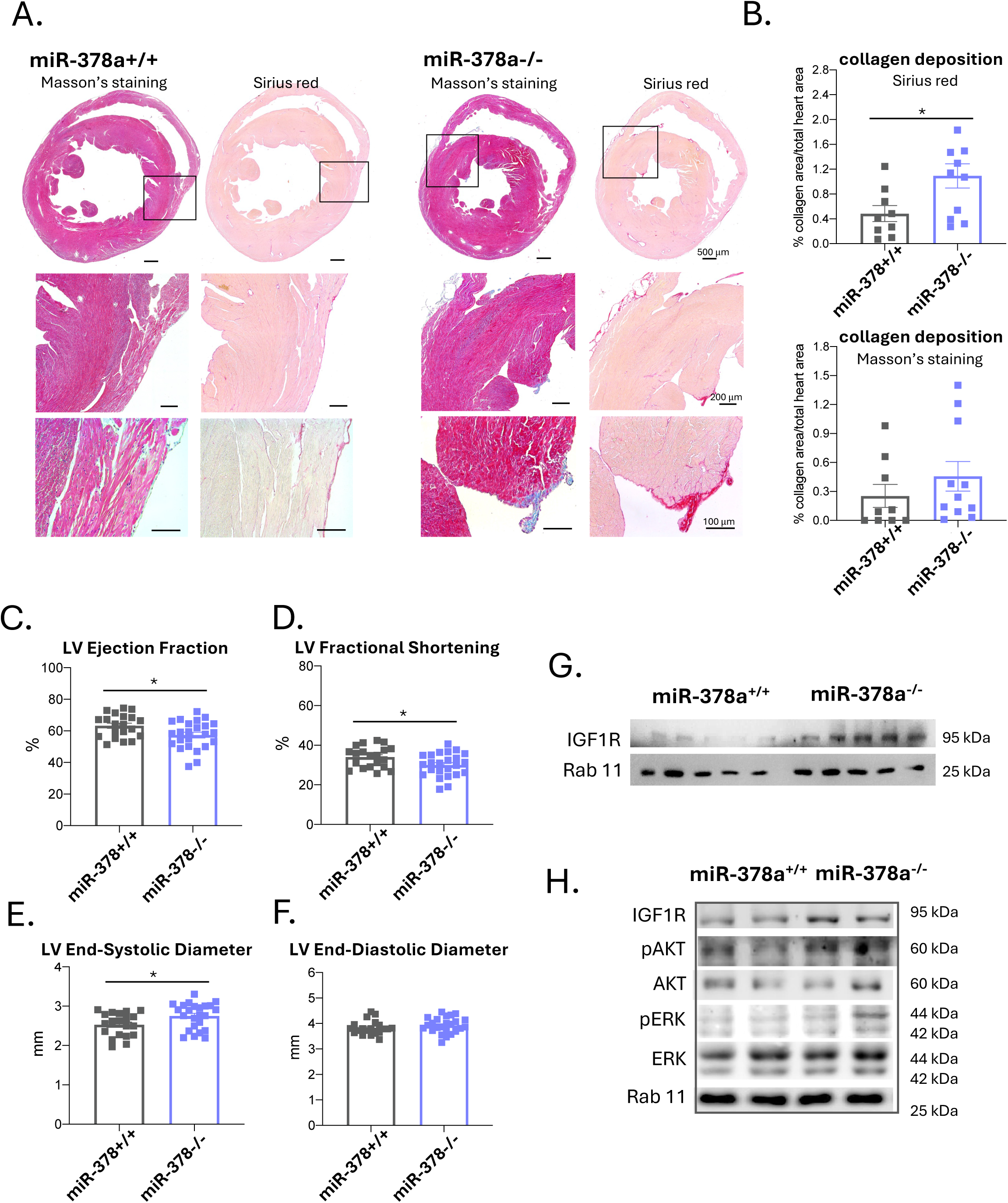
**A-B. (A)** Representative cardiac sections and (**B**) Semiquantitative analysis of collagen deposition in cardiac sections from 11 – 12-week-old miR-378a+/+ and miR-378a-/-mice (n=9-11). *p<0.05. **G-J.** Left ventricle (LV) Ejection Fraction (**G.)**, LV Fractional Shortening (**H.)**, LV End-Systolic volume (**I.**) and LV End-Diastolic volume (**J.**) in 10 – 13-week-old miR-378a+/+ (n=20) and miR-378-/- mice (n=24) *p<0.05, unpaired two-tailed Student’s t-test. **G.** Western blot analysis of IGF1R pathway in heart lysates isolated from (**G**)11 - 12-week-old and (**H**) 6-week-old miR-378a+/+ and miR-378a-/- mice (n=5, n=2, respectively). Rab11 served as a protein loading control.

Interestingly, in 17-month-old animals EF and FS parameters were similar in both genotypes (Supplementary Fig. 2A, B), while end-systolic volume and diameter (Supplementary Fig. 2C, E), end-diastolic volume and diameter (Supplementary Fig. 2D, F) as well as SV (Supplementary Fig. 2G) were significantly decreased in miR-378a-deficient mice. Thus, at both time points, miR-378a knockout affected heart function. However, the impact on different cardiac parameters in younger and older animals suggests that the physiological role of miR-378a in the myocardium may change with age.

IGFR signaling has previously been implicated in mediating the effects of miR-378a on cardiac hypertrophy (7). Consistently, IGF1R protein levels were upregulated in miR-378a−/− heart extracts from 11–12-week-old mice (Fig. 1G). Importantly, already in 6-week-old mice, key components of this pathway, including IGF1R, phospho-AKT (pAKT) and phospho-ERK1/2 (pERK), were elevated in miR-378a-/- hearts compared with wild-type controls (Fig. 1H).

### 3.2. Integrated global transcriptomic and proteomic analyses of miR-378a-deficient hiPSC-CM and murine hearts

Echocardiographic analysis of miR-378a-/- mice confirmed that lack of miR-378a affects heart function even under basal, non-pathological conditions. To gain further insight into the molecular processes regulated by this miRNA in cardiomyocytes, both murine and human, we performed integrated analysis of the transcriptomic and proteomic data generated from control and miR-378a-deficient hiPSC-CM as well as proteomic data generated from miR-378a+/+ and miR-378a-/- murine hearts.

Given the complexity of the intended approach and the limitations in the available cell lines, in the first step we performed transcriptomic analysis of hiPSC-CM obtained from control and miR-378a-deficient hiPSC1 line, previously described by us (8). In particular, four independent clones of hiPSC1 with confirmed deletion of *MIR378A* locus (clone 16, 23, 44 and 48) (8), together with the isogenic control cells, were subjected to cardiac differentiation.

After confirming efficient hiPSC-CM differentiation (>80% cardiac troponin T (cTnT)-positive cells; data not shown), RNA samples were collected for global transcriptomic analysis. This initial step revealed considerable heterogeneity even between distinct clones derived from the same parental cell line. Following the removal of the clone effect, the analysis revealed 94 significantly downregulated transcripts in the miR-378a-deficient hiPSC-CM lines (adjusted p-value < 0.05). Of these, 17 transcripts showed more than 4-fold change in expression (|log_2_ fold-change| > 2). Additionally, 39 transcripts were found to be significantly upregulated, 14 of which showed more than 4-fold change in expression (Supplementary Fig. 3A, Supplementary Table 1). Gene ontology (GO) analysis of the genes with significantly altered expression revealed 85 over-represented GO-terms (adjusted p-value < 0.05). For downregulated genes, 58 GO-terms belonged to Biological Processes (BP), 24 to Cellular Components (CC) and 2 to Molecular Functions (MF) (Supplementary Fig. 3B, Supplementary Table 2). Upregulated genes resulted in only one overrepresented GO-term: I band (belonging to CC, Supplementary Fig. 3B). Many of the depicted processes were related to muscle contraction and regulation of the action potential (Supplementary Fig. 3B). Additionally, using the same criteria 15 significantly over-represented KEGG pathways were detected for downregulated genes, among which were those involved in dilated and hypertrophic cardiomyopathy (Supplementary Fig. 3C, Supplementary Table 3). No over-represented KEGG pathways were detected for upregulated genes.

Taking into consideration that miRNAs predominantly regulate gene expression on post-transcriptional level, in the next step we performed global proteomic analysis of control and miR-378a-deficient hiPSC-CM. Based on our previous study (12) in which we observed strong effect of donor’s genetic background on the transcriptome and proteome of hiPSC-CM, to decrease the risk of observing single donor-related changes, we first decided to generate novel hiPSC lines with CRISPR/Cas9-mediated deletion of *MIR378A* locus. Additionally, to minimize the impact of possible off-target activity of CRISPR/Cas9 system applied previously (8), we designed new sgRNA sequences targeting 5’ upstream and 3’ downstream of *MIR378A* gene and delivered them together with Cas9 nuclease to the hiPSC2 (generated from PBMC of male donor) and hiPSC3 (generated from PBMC of female donor) lines using the same approach as for hiPSC1 line (8). Successful deletion of *MIR378A* in selected hiPSC2 and hiPSC3 clones was validated using PCR-based genotyping (data not shown) and lack of miR-378a-3p and miR-378a-5p strand expression was further confirmed before (Supplementary Fig. 4A, B) and after cardiac differentiation using qPCR method (Supplementary Fig 4C, D). Additionally, miR-378a-deficient hiPSC2 and hiPSC3 lines retained expression of pluripotency markers (Supplementary Fig. 4E, F) and PGC1β, the latter indicating that CRISPR/Cas9 approach engaged in our study did not inactivate the gene within which the *MIR378A* locus is located (data not shown). Eventually, for the global proteomic readout two miR-378a knock-out clones (44 and 48, presenting the most consistent transcriptome) from hiPSC1 line as well as single miR-378a knock-out clone from hiPSC2 and hiPSC3 lines were subjected to cardiac differentiation, total protein isolation and mass spectrometric measurement. Separate analysis of obtained data, however, revealed no significantly altered proteins in miR-378a-deficient cardiomyocytes in comparison to their control counterparts, possibly due to strong impact of donor’s genetic background on global proteome of hiPSC-CM (Supplementary Fig. 5).

In parallel, we decided to complement our studies with global proteomic analysis of miR-378a+/+ and miR-378a-/- hearts isolated from 10 – 11-week-old animals. For that purpose the entire organs were perfused with heparinized saline (0.5 IU/ml) and 30 mM KCl, snap-frozen in liquid nitrogen, subjected to protein extraction with 1% SDS in 0.1 M Tris-HCl and eventually directed to mass spectrometric measurement of the protein content.

The proteomic differential expression analysis of hiPSC-CM revealed no proteins with significantly altered expression (Supplementary Figure 5), while the analysis of murine hearts showed only one slightly over-expressed protein (Q8JZQ2, Supplementary Figure 6).

To obtain an overall picture from the transcriptomic and proteomic data, we combined all omics datasets, including transcriptomic and proteomic data from hiPSC-CM and proteomic data from murine hearts. We then compared the direction of expression changes for all genes with known one-to-one orthologues between humans and mice according to the Ensembl database, regardless of statistical significance in the individual datasets.

For subsequent functional analysis, we used all genes showing a common pattern across all three datasets, i.e. consistent increase (364 genes) or decrease (266 genes) in expression. Such an approach revealed 46 over-represented GO-terms for persistently over-expressed genes (adjusted p-value < 0.05, Fig. 2A, Supplementary Table 4), encompassing 27 BPs, 6 MFs and 12 CCs. Among them many are involved in protein-related catabolic and biosynthetic processes including translation regulatory activity and post-transcriptional gene expression. Additionally, 46 altered KEGG pathways (21 over-expressed and 25 under-expressed) (Fig. 2B, Supplementary Table 5) were detected with a strong indication of disturbed metabolism in the absence of miR-378a.

**Figure 2.**
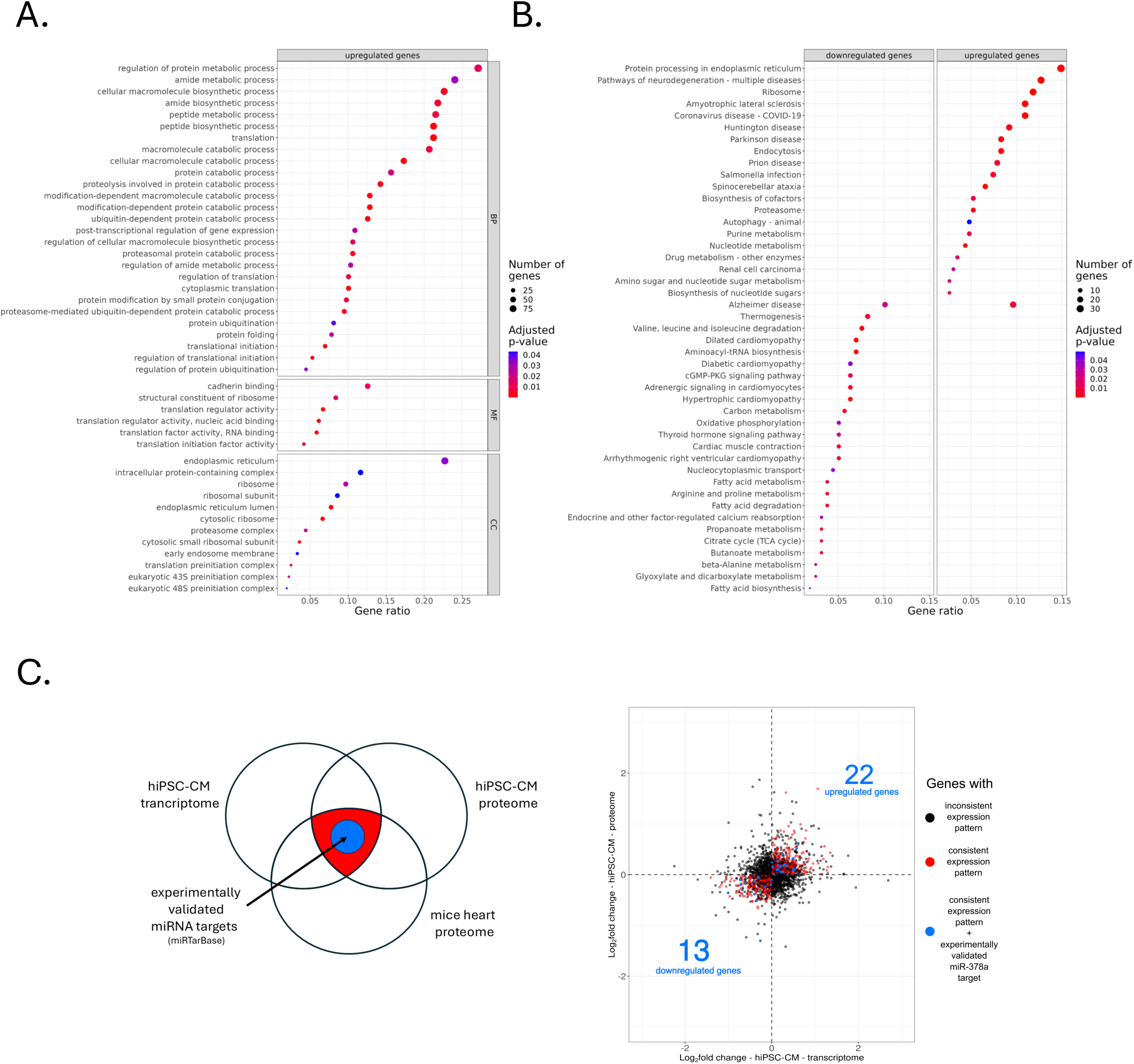
**A.** Dot plot representing GO-terms over-representation analysis of combined transcriptomic and proteomic dataset from control and miR-378aKO hiPSC-CM as well as proteomic dataset from miR-378a+/+ and miR-378a-/- hearts of 11-12-week-old mice. Only genes with a consistent direction of expression change across all three datasets were included in the analysis. The plot only shows the analysis of GO-terms for consistently upregulated genes, as no significant over-represented GO-terms were found for consistently downregulated genes. BP – biological processes, MF – molecular function, CC – cellular components. **B.** Dot plot of KEGG pathways over-representation analysis for combined transcriptomic and proteomic dataset from control and miR-378aKO hiPSC-CM as well as proteomic dataset from miR-378a+/+ and miR-378a-/- hearts of 11-12-week-old mice. Only genes with a consistent direction of expression change across all three datasets were included in the analysis. **C.** The results of combined analysis of control and miR-378aKO hiPSC-CM transcriptomic and proteomic data as well as proteomic data from miR-378a+/+ and miR-378a-/- hearts integrated with experimentally validated miR-378a targets accessed from miRTarBase.

Finally, we narrowed down the set of genes incorporating the miRTarBase database (13), consisting of experimentally validated miRNA targets. We found that 22 of consistently upregulated and 13 of consistently downregulated genes in miR-378a-deficient samples corresponded to experimentally validated miR-378a targets (Fig. 2C). Interestingly, among them were factors with important function in cardiomyocyte physiology including Yin Yang 1 (YY1) (14) and cholinergic receptor muscarinic 2 (CHRM2) (15) (Supplementary Fig. 7, highlighted in red).

### 3.3. miR-378a knockout affects metabolism of human cardiomyocytes

Among all altered pathways revealed in our integrated analysis of the omics data from murine hearts and hiPSC-CM, we noticed the possible effect of miR-378a on cardiac metabolism. This is in line with the previously described function of the investigated miRNA in maintaining systemic metabolic homeostasis (2, 3). Interestingly, the role of miR-378a in regulating metabolic processes in murine and human cardiomyocytes has not been thoroughly studied so far. To fill this gap, we focused our research on hiPSC-CM which better reflects the human cardiomyocyte physiology. In the first step, the response of control (CTR) and miR-378a-knockout (miR-378aKO) hiPSC1-CM to inhibition of mitochondrial respiratory chain subunits was verified using the Seahorse XF Mito Stress method. Obtained results indicate that miR-378a knockout in hiPSC-CM reduces the oxygen consumption rate (OCR), particularly after the application of the FCCP (p-trifluoromethoxyphenyl carbonyl cyanide hydrazone), which uncouples the mitochondrial electron chain (Fig. 3A). Additionally, performed analysis revealed that miR-378aKO hiPSC1-CM demonstrate reduced basal respiration (Fig. 3B), ATP production (Fig. 3C), maximal respiration (Fig. 3D), non-mitochondrial respiration (Fig. 3E) and spare respiratory capacity (Fig. 3F) compared to their control counterparts. To further evaluate the underlying mechanism of such changes we performed PCR-mediated quantification of mitochondrial to genomic DNA ratio and observed decreased mitochondrial content in miR-378aKO hiPSC1-CM (Fig. 3G). In parallel, transmission electron microscopy (TEM) revealed that miR-378a-deficiency was associated with the disrupted morphology of mitochondria which appeared to be swollen and characterized by abnormal mitochondrial crista structure (Fig. 3H).

**Figure 3.**
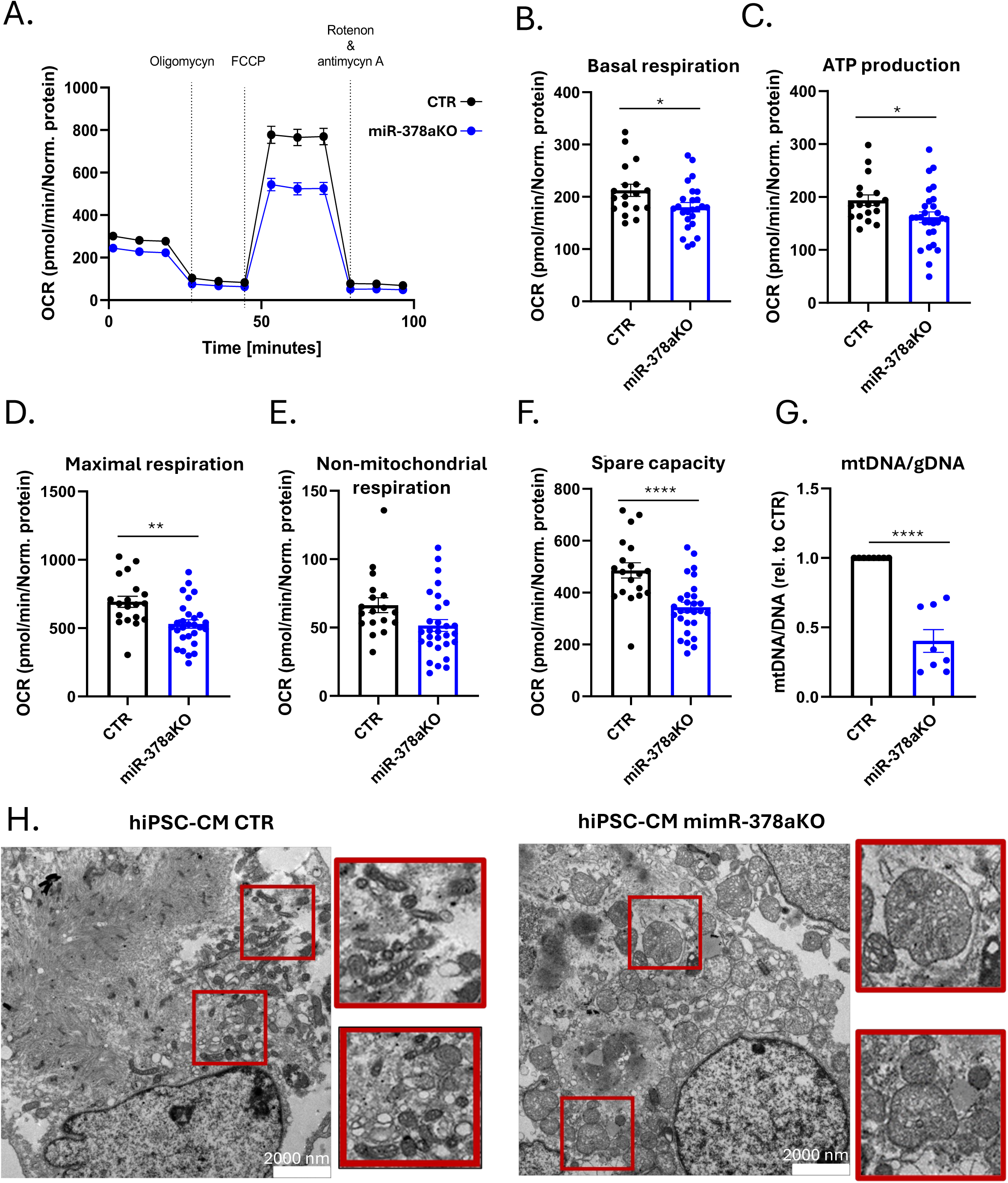
**A.** Oxygen consumption rate (OCR) in control (CTR) and miR-378aKO hiPSC-CM analysed by Seahorse XF Mito Stress assay. FCCP - trifluoromethoxy carbonylcyanide phenylhydrazone, carbonyl cyanide 4- (trifluoromethoxy)phenylhydrazone. **B – F.** Basal respiration (**B.**), ATP production (**C.**), maximal respiration (**D.**), non-mitochondrial respiration (**E.**) and spare respiratory capacity (spare capacity) (**F.**) in control (CTR) and miR-378aKO hiPSC1-CM (n=3), *p<0.05, **p<0.01, ****p<0.0001, unpaired two-tailed Student’s t-test. **G.** qPCR analysis of mitochondrial (mtDNA) to genomic (gDNA) ratio in control (CTR) and miR-378aKO hiPSC-CM. n=3, ****p<0.0001, unpaired two-tailed Student’s t-test. **H.** Representative TEM images of control (CTR) (left panel) and miR-378aKO hiPSC-CM (right panel). Red boxes magnify selected mitochondria, scale bar = 2 µm.

### 3.4. miR-378a regulates mitophagy in hiPSC-CM

Mitochondrial content strongly relies on the proper balance between mitochondrial biosynthesis and mitophagy. Therefore, we focused our next steps on investigating these processes in control and miR-378a-KO hiPSC-CM. Importantly, TEM data suggested possible alterations in mitochondrial morphology consistent with increased mitophagy in the latter cells. To further validate this observation, cells were stimulated with 20 μM CCCP, a compound that uncouples the electron transport chain by depolarizing the mitochondrial membrane upon which the level of mitophagy-related proteins was examined. Parkin was elevated in miR-378aKO hiPSC1-CM both before and after mitophagy induction in comparison to their control counterparts, regardless of the CCCP-mediated decrease in the level of this protein in both genotypes (Fig. 4A). Parkin is essential for the induction of mitophagy, marking defective mitochondria (16). Receptors such as p62 and OPTN are then involved in this process. In our experiments, higher level of p62 was observed in miR-378aKO hiPSC1-CM both before and after mitophagy induction (Fig. 4A), whereas OPTN levels decreased after CCCP treatment only in control cells, remaining unchanged in the miR-378aKO hiPSC1-CM group (Fig. 4A). Interestingly, the level of lipidated LC3 (LC3II) was lower in miR-378aKO hiPSC1-CM after CCCP treatment compared to control cardiomyocytes (Fig. 4A). Further flow cytometric analysis using autolysosome-specific probe indicated that upon CCCP treatment, miR-378aKO hiPSC1-CM demonstrated decreased formation of these autophagic vacuoles (Fig. 4B).

**Figure 4.**
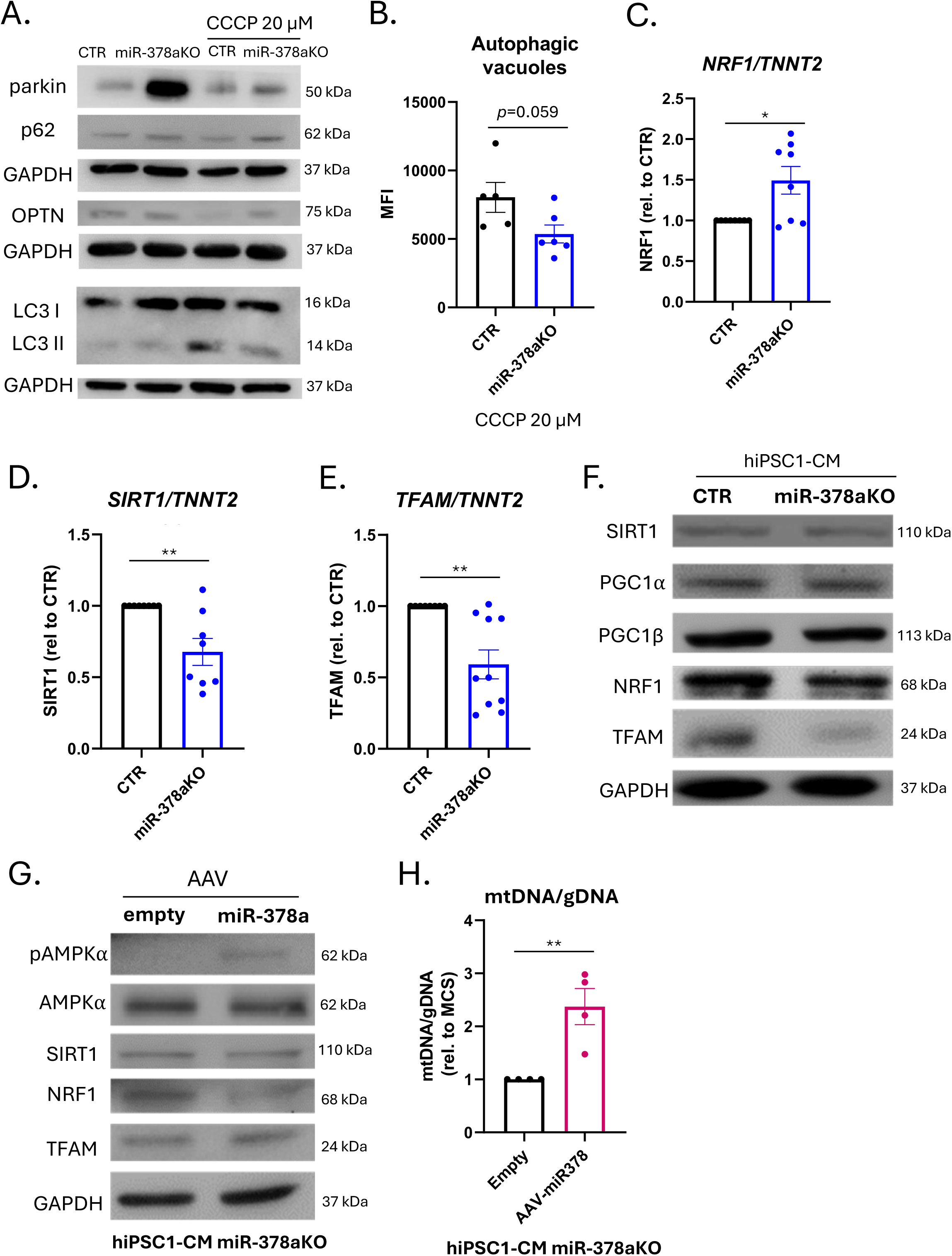
**A.** Representative Western blot for the detection of mitophagy-related proteins: parkin, p62, OPTN, LC3 I and II in control (CTR) and miR-378aKO hiPSC-CM, non-treated and treated with 20 µM CCCP. GAPDH served as a protein loading control. n=6 (two cell lines were used, hiPSC-CM1 and hiPSC-CM2, for each 3 replicates were performed). **B.** Detection of autophagic vacuoles in control (CTR) and miR-378aKO hiPSC1-CM cells after mitophagy induction with 20μM CCCP using a specific probe and flow cytometry, N=6 ± SEM (two cell lines were used, hiPSC1-CM and hiPSC2-CM, for each 3 replicates were performed), *p<0.05, unpaired two-tailed Student’s t-test. **C – E.** qPCR analysis of NRF1 (**C.**), SIRT1 (**D.**) and TFAM (**E.**) in control (CTR) and miR-378aKO hiPSC1-CM. TNNT2 was used as a reference control. n=3, *p<0.05, unpaired two-tailed Student’s t-test. **F.** Representative Western blot for the detection of mitochondrial biosynthesis-involved protein: SIRT1, PGC1α, PGC1β, NRF1 and TFAM in control (CTR) and miR-378aKO hiPSC1-CM. GAPDH served as a protein loading control. **G.** Representative Western blot for the detection of mitochondrial biosynthesis-involved protein: phosphoAMPKα (pAMPKα), total AMPKα, SIRT1, NRF1 and TFAM in miR-378aKO hiPSC-CM transduced with AAV6-empty (empty) and AAV6-miR-378a (miR-378a). GAPDH served as a protein loading control. **H.** qPCR analysis of mitochondrial (mtDNA) to genomic (gDNA) ratio in miR-378aKO hiPSC-CM transduced with AAV6-empty (empty) and AAV6-miR-378a (miR-378a). n=4, **p<0.01, unpaired two-tailed Student’s t-test.

### 3.5. miR-378a regulates mitochondrial biogenesis in hiPSC-CM

Observed changes in miR-378a-KO cardiomyocytes indicated disturbed mitophagy-mediated turnover of mitochondria in these cells. Thus, in the next step we examined the counteractive process of mitochondrial biogenesis. The signaling pathway regulating this process is activated by an increase in the AMP/ATP and NAD+/NADH ratios, which translates into the activity of AMPK and SIRT1, respectively. These proteins mediate nuclear translocation of PGC1α, where it cooperates with transcription factor NRF1 to induce the expression of genes involved in mitochondrial biosynthesis, including *TFAM*, *GABPA* (encoding nuclear respiratory factor 2 (NRF2)), *CREB1*, *SIRT1*, and *PPRGC1A* (encoding PGC1α). TFAM is, in turn, translocated to mitochondria, where it mediates mtDNA replication (17). First, we observed no differences in *NRF1* expression (Fig. 4C) in miR-378aKO hiPSC1-CM in comparison to their control counterparts while *SIRT1* (Fig. 4D) and *TFAM* (Fig. 4E) were significantly downregulated in these cells. Importantly, similar pattern was also observed in miR-378a-KO hiPSC2-CM (generated from hiPSC2 line), with decrease in TFAM, and SIRT1, although the change in *SIRT1* did not reach statistical significance (Supplementary Fig. 8A-C). The decrease in TFAM level was also visible in Western blot analysis (Fig. 4F) and representative immunofluorescent staining (Supplementary Fig. 8D). A slight decrease in NRF1 protein level was also observed in miR-378aKO hiPSC1-CM, while, interestingly, no differences in SIRT1 as well as PGC1 α/β levels were detected between both genotypes (Fig. 4F) indicating a possible compensatory mechanism preventing downregulation of SIRT1 protein level. To verify the extent to which the observed changes were dependent on miR-378a, the expression of this miRNA was restored in miR-378aKO hiPSC1-CM using AAV6 vectors which resulted in the upregulation of pAMPKα and TFAM (Fig. 4G) and downregulation of NRF1 (Fig. 4G), while no effect was observed for SIRT1 and total AMPKα (Fig. 4G). Importantly, transduction of miR-378a-KO cardiomyocytes with AAV6-miR-378a vectors was associated with increased mtDNA/gDNA ratio (Fig. 4H) indicating positive effect on mitochondrial content in these cells. Successful restoration of miR-378a-3p and miR-378a-5p expression in miR-378aKO hiPSC1-CM was confirmed with qPCR method (Supplementary Fig. 8E).

### 3.6. miR-378a regulates glucose uptake and glycogen synthesis

The results of Seahorse XF Mito Stress analysis prompted us predominantly to investigate mitochondrial homeostasis in mir-378a-KO hiPSC-CM, however, the same assay additionally revealed that miR-378aKO hiPSC1-CM acidify the medium to a lesser extent indicating the possible effect of miR-378a knockout on glycolysis (Supplementary Fig. 9A). Of note, miRNA-378a has been already described to regulate this process in cancer cells (18). Thus, our next step was to examine the glucose processing pathways in human cardiomyocytes. Performed experiments demonstrated a tendency for miR-378aKO hiPSC1-CM to increase glucose uptake in comparison to their control counterparts (p=0.07) (Fig. 5A). Interestingly, this was accompanied with significantly decreased hexokinase (Fig. 5B) and LDH (Fig. 5C) activities as well as elevated levels of the glucose transporter GLUT1 (Fig. 5D) and the γ-subunit of AMPK, responsible for sensing the amount of AMP relative to ATP in the cells (Fig. 5D). Of note, after restoration of miR-378a expression in miR-378aKO hiPSC1-CM using AAV6-miR-378a vectors, GLUT1 level decreased substantially (Fig. 5E). In parallel, PAS staining demonstrated a decrease in glycogen accumulation in miR-378aKO hiPSC1-CM in comparison to the control group (Fig. 5F), despite increased glucose uptake in these cells. In line with this observation, the levels of active and basal forms of GSK3α/β were elevated in miR-378aKO hiPSC1-CM (Fig. 5G), and restoration of miR-378a expression slightly reversed this effect (Fig. 5H). Similar results in terms of GLUT1, AMPKγ and GSK3α/β levels were obtained in miR-378aKO hiPSC2-CM (Supplementary Fig. 9B).

**Figure 5.**
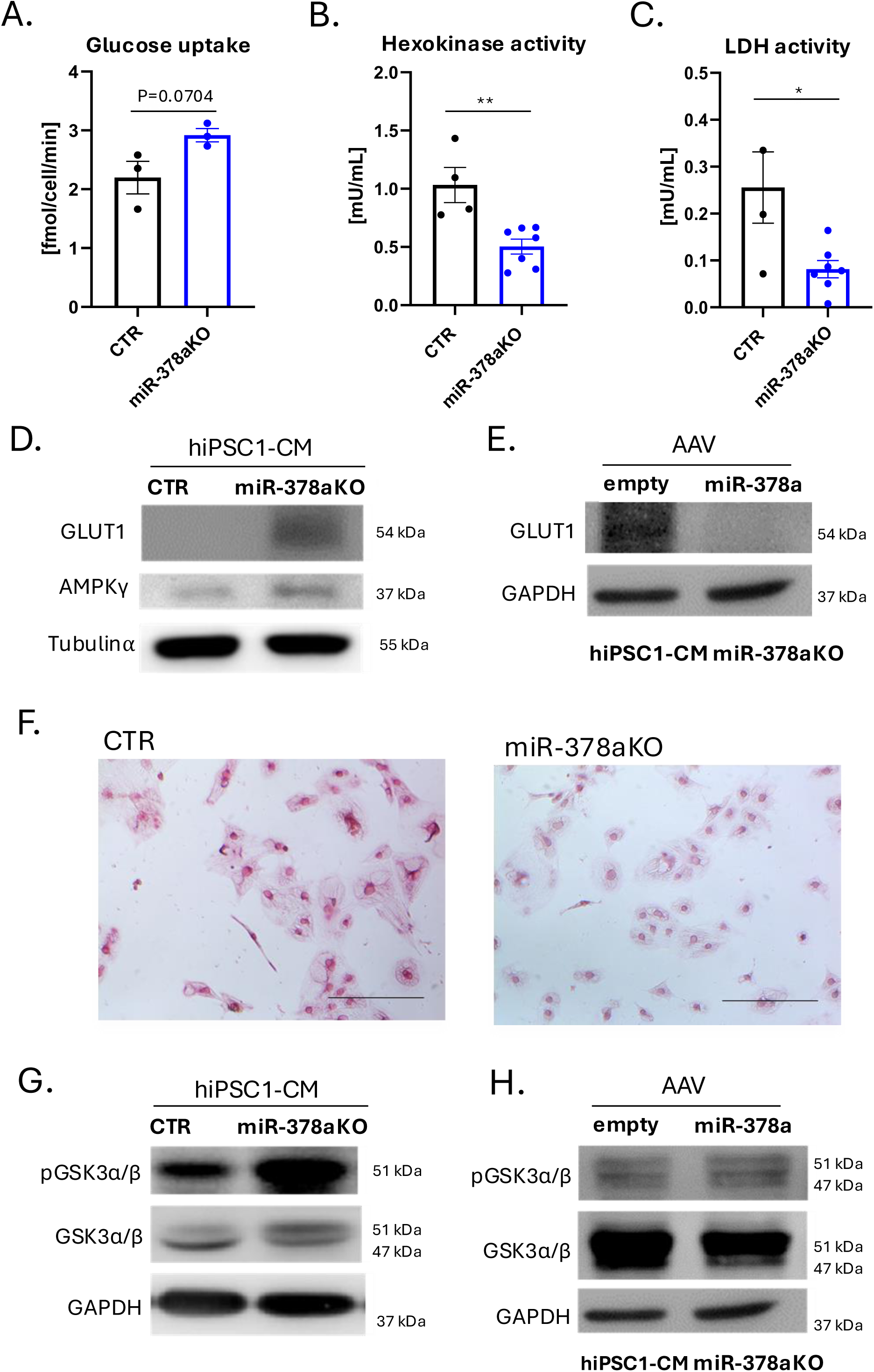
**A – C.** Glucose uptake (**A.**), hexokinase activity (**B.**) and LDH activity (**C.**) in control (CTR) and miR-378aKO hiPSC1-CM. n=4, *p<0.05, unpaired two-tailed Student’s t-test. **D.** Representative Western blot for GLUT1 in control (CTR) and miR-378aKO hiPSC1-CM. β Tubulin served as a protein loading control. **E.** Representative Western blot for GLUT1 in miR-378aKO hiPSC1-CM transduced with AAV6-empty (empty) and AAV6-miR-378a (miR-378a). GAPDH served as a protein loading control. **F.** Representative images of control (CTR) and miR-378aKO hiPSC1-CM staining using the glycogen-detecting PAS method, images were taken at 200x magnification, scale bar = 50 µm. **G.** Representative Western blot for phosphoGSK3α/β on Tyr206 (GSK3α/β p Tyr206) and total GSK3α/β in control (CTR) and miR-378aKO hiPSC1-CM. GAPDH served as a protein loading control. **H.** Representative Western blot for for phosphoGSK3α/β on Tyr206 (GSK3α/β p Tyr206) and total GSK3α/β in miR-378aKO hiPSC1-CM transduced with AAV6-empty (empty) and AAV6-miR-378a (miR-378a). GAPDH served as a protein loading control.

### 3.7. miR-378a regulates angiogenic activity of hiPSC-CM

Having observed that the knockout of miR-378a disturbs cardiomyocyte metabolism on different levels, we next investigated whether it may also affect properties more directly related to therapeutic potential of hiPSC-CM in cardiac cell therapies. Particularly, our previous study highlighted pro-angiogenic activity of miR-378a in skeletal muscles manifesting, among others as decreased number of CD31-positive blood vessels and arterioles in gastrocnemius muscle of miR-378a-/- mice (4). While studying the role of miR-378a in streptozotocin-induced diabetic cardiomyopathy we also observed the tendency towards reduced number of α smooth muscle actin (αSMA)-positive vessels in vehicle-treated miR-378a-/- hearts in comparison to their wild type counterparts (10). In the current study we extended the aforementioned analysis and performed the immunohistochemical examination of αSMA-, CD31- and lectin-positive vessels in 11 – 12-week-old miR-378a+/+ and miR-378a-/- hearts, however, the observed decrease in miR-378a-KO specimen did not reach statistical significance (Supplementary Fig. 1E). Nevertheless, we decided to complement this investigation with the evaluation of pro-angiogenic activity of control and miR-378aKO hiPSC-CM. For that purpose, matrigel tube formation assay was performed using human aortic endothelial cells (HAEC line) seeded on thick layer of matrigel. Importantly, tube formation was significantly impaired when cells were cultured in miR-378aKO hiPSC1-CM-conditioned medium (Fig. 6A). Quantitative analysis 24 hours after seeding demonstrated significantly decreased total tube length (Fig. 6B), number of nodes (Fig. 6C), branches (Fig. 6D) and junctions (Fig. 6E) in this group in comparison to the control counterparts. Counterintuitively, however, miR-378a-KO hiPSC-CM secreted more VEGF than control cells (Fig. 6F) indicating that the concomitant production anti-angiogenic factors may overwhelm the effect of VEGF or the level of VEGF may be too high for proper formation of tubular structures. Finally, we evaluated the angiogenic potential of hiPSC1-CM in 3D cell culture setting using idenTx 3 microfluidic chip which was composed of fibrin hydrogel-filled middle channel enabling cell migration and two lateral channels: one seeded with HAEC and the opposite seeded with either control or miR-378aKO hiPSC-CM. Surprisingly, instead of observing migration of endothelial cells toward cardiomyocytes, we noticed propagation of control hiPSC-CM into the hydrogel space which was strongly inhibited in miR-378a-KO cells (Supplementary Fig. 10). This might be in line, however, with the transcriptomic data which revealed under-expression of several biological processes related to cell-cell and cell-matrix adhesion as well as actin filament-based movement in miR-378aKO hiPSC-CM (Supplementary Fig. 3C).

**Figure 6.**
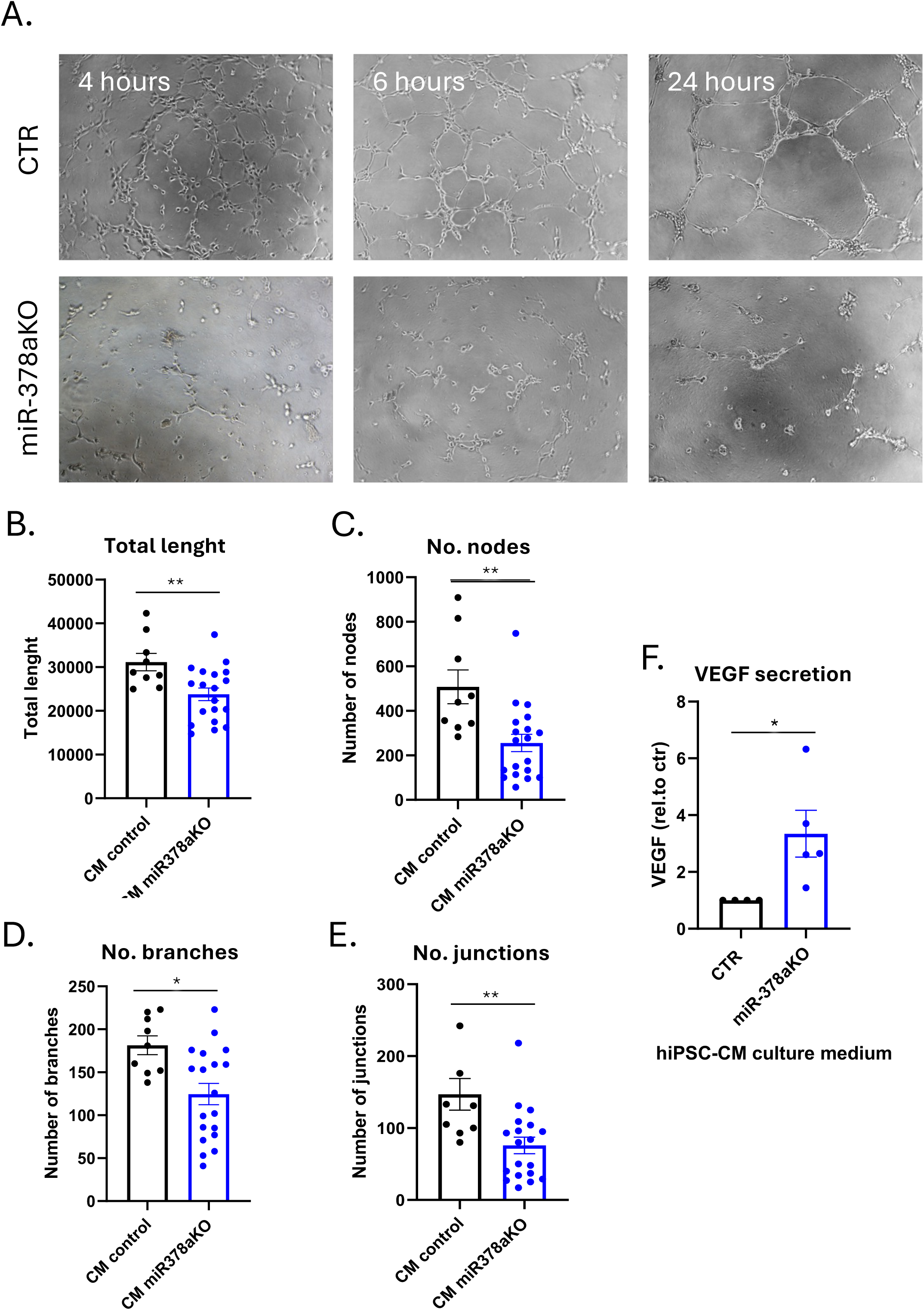
**A.** Representative images of HEAC cells 4, 6 and 24 hours after seeding on matrigel in conditioned media collected from control (CTR) and miR-378aKO hiPSC-CM. **B – E.** Total length (**B.**), number of nodes (**C.**), number of branches (**D.**) and number of junctions (**E.**) in tubes formed by HAEC 24 hours after seeding on matrigel in conditioned media collected from control (CM control) and miR-378aKO (CM miR-378aKO) hiPSC-CM. n=3, *p<0.05, **p<0.01, unpaired two-tailed Student’s t-test. **F.** ELISA-based analysis of VEGF secretion by control (CTR) and miR-378aKO hiPSC-CM. N=3, *p<0.05, unpaired two-tailed Student’s t-test.

### 3.8. miR -378a overexpression does not improve the therapeutic potential of hiPSC-CM-based cell therapy in murine model of acute myocardial infarction

Taking into consideration that miR-378a knockout disturbs metabolic homeostasis and pro-angiogenic activity of hiPSC-CM, in the last step we hypothesized that its overexpression may increase the therapeutic potential of these cells. To verify this concept, we transduced control hiPSC-CM with AAV6-Luc encoding luciferase for *in vivo* bioimaging (Supplementary Fig. 11A) as well as either with AAV6-empty or AAV6-miR-378a vectors (level of miR-378a-3p and miR-378a-5p overexpression presented in Supplementary Fig. 11B) and administered such cardiomyocytes into the hearts of NOD-SCID mice subjected to myocardial infarction. IVIS-based monitoring of luciferase activity upon peritoneal luciferin injection revealed selected animals in which the bioluminescence was present 2 weeks and 6 weeks after cell delivery (Supplementary Fig. 11C). In the latter time point the presence of both hiPSC-CM AAV-empty and hiPSC-CM AAV-miR-378a, manifested as luciferase activity, was particularly visible in isolated hearts at the 6-week endpoint of the study (Supplementary Fig. 11D). Importantly, echocardiographic analysis of heart function 2, 4 and 6 weeks after cell delivery demonstrated that administration of both hiPSC-CM AAV-empty and AAV-miR-378a significantly improved left ventricle (LV) EF (Fig. 7A) and FS (Fig. 7B), however, at 4-week time point only and with no additive effect of mir-378a overexpression. Similarly, LV end-systolic (Fig. 7C) and LV end-diastolic (Fig. 7D) diameters were significantly decreased upon administration of both types of hiPSC-CM at 4-week time point with no additional effect of miR-378a overexpression. Taking into consideration that hiPSC-CM AAV-miR-378a did not provide better therapeutic outcome compared to their control counterparts in the last set of experiments we examined whether cell delivery induced any molecular changes in the infarcted heart milieu. Performed qPCR analysis revealed that MI induction resulted in decreased cardiac expression of miR-378a-3p and miR-378a-5p (Supplementary Fig. 12A, B), which was upregulated in both hiPSC-CM-treated groups with stronger effect of hiPSC-CM AAV-miR-378a, possibly indicating detection of human miR-378a in the analysis. Interestingly, administration of hiPSC-CM AAV-miR378a resulted in significantly increased expression of murine VEGFA on mRNA level compared to the vehicle- and control cell-treated hearts (Fig. 7E), however, the corresponding increase of VEGFA protein in this group did not reach statistical significance (Fig. 7F). In parallel, we observed the tendency to similar pattern of *Kdr* (encoding VEGFR2) upregulation (hiPSC-CM AAV-miR-378a vs vehicle and control cell groups, p=0.0544 and p=0.0501, respectively) (Supplementary Fig. 12C). Finally, hearts subjected to MI demonstrated the increase in murine *Cxcr4* transcript level in comparison to sham control, which reached statistical significance in hiPSC-CM AAV-miR-378a-treated animals (Supplementary Fig. 12D). Otherwise, no significant changes were observed in the expression of *Cxcl12* (encoding SDF-1 chemokine) (Supplementary Fig. 12E), *Fgf1* (Supplementary Fig. 12F), *Col1a1* (encoding collagen type 1α) (Supplementary Fig. 12G), *Fap* (encoding fibroblast activation protein α) (Supplementary Fig. 12H) and *Fn1* (encoding fibronectin-1) (Supplementary Fig. 12I). In summary, although miR-378a overexpression did not improve heart function upon hiPSC-CM delivery in acute MI model, it induced distinct molecular changes in the infarcted myocardium. Further studies, including evaluation of longer time points and administration of higher cell number, are needed to better assess whether such changes may eventually result in a better functional outcome.

**Figure 7.**
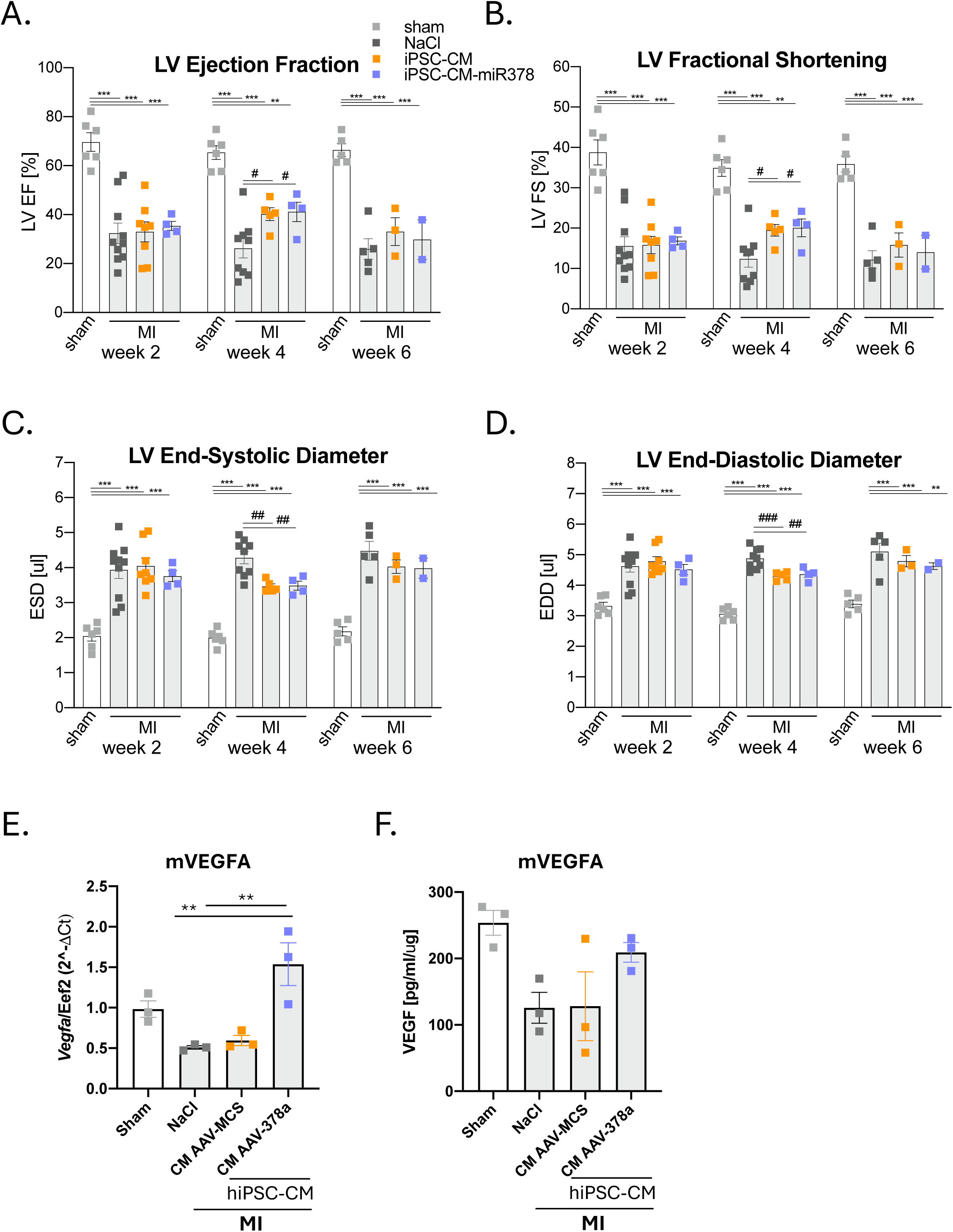
**A – D.** LV Ejection Fraction (**A.**), LV Fractional Shortening (**B.**), LV End-Systolic Volume (**C.**) and LV End-Diastolic Volume (**D.**) two, four and 6 weeks after MI induction in mice treated with vehicle (NaCl), hiPSC-CM transduced with AAV6-empty vectors (iPSC-CM) and hiPSC-CM transduced with AAV6-miR-378a vectors (iPSC-CM-miR-378a). Mice subjected to sham procedure served as a control. n=2-10, *p<0.05, **p<0.01, ***p<0.001 sham vs MI groups; #p<0.05, ##p<0.01, ###p<0.001 NaCl vs hiPSC-CM-treated groups, two-way ANOVA, followed by Tukey’s post hoc test + unpaired two-tailed t-test (Welch’s correction where appropriate) **E.** qPCR analysis of Vegfa (encoding murine VEGFA, mVEGFA) expression in heart lysates collected 6 weeks after MI induction from mice treated with vehicle (NaCl), hiPSC-CM transduced with AAV6-empty vectors (AAV-empty) and hiPSC-CM transduced with AAV6-miR-378a vectors (AAV-miR-378a). Mice subjected to sham procedure served as a control. n=3 animals in each group, **p<0.01, one-way ANOVA, followed by Tukey’s post hoc test. **F.** ELISA-based analysis of murine VEGF (mVEGF) protein level in heart lysates collected 6 weeks after MI induction from mice treated with vehicle (NaCl), hiPSC-CM transduced with AAV6-empty vectors (AAV-empty) and hiPSC-CM transduced with AAV6-miR-378a vectors (AAV-miR-378a). Mice subjected to sham procedure served as a control. n=3 animals in each group.

## 4. Discussion

This study provides comprehensive insights into the role of miR-378a in the regulation of cardiomyocyte physiology and metabolism under basal, non-pathological conditions and in response to injury, in post-MI cell therapy approach in mice. Using a combination of murine and human models, as well as multi-omics, we demonstrate that miR-378a plays a multifaceted role in maintaining myocardial function, mitochondrial homeostasis, glucose metabolism, and cardiac angiogenic signaling. Integrated transcriptomic and proteomic analyses in miR-378a knockout and control mice, as well as hiPSC-CM of both genotypes, reveal miR-378a influence on pathways related to translation, metabolism, and cardiomyopathy-associated signaling. *In vivo*, miR-378a knockout in mice promotes myocardial fibrosis, alters hypertrophic signaling, and impairs cardiac function, with age-specific differences in affected parameters. Conversely, following myocardial infarction, AAV-mediated miR-378a overexpression in hiPSC-CM did not further enhance the already beneficial effect of these cells in NOD-SCID mice. Our findings suggest that while miR-378a is critical for maintaining cardiomyocyte metabolic homeostasis and structural integrity, its therapeutic overexpression post-MI may offer limited additional benefit due to the dominance of alternative repair pathways.

Until now, the research on the role of miR-378a in the heart mainly concerned its activity in pathological processes such as chronic hypertension (*in vivo*), hypertrophy (*in vitro, in vivo*), and the response of cardiomyocytes and fibroblasts to hypoxia (*in vitro*) [7,19–21]. Our echocardiographic analyses revealed that miR-378a knockout in mice leads to significant impairment of cardiac function, even in the absence of pathological stress. Younger miR-378a knockout animals displayed signs of early ventricular dysfunction (reduced EF and FS), while older mice exhibited broader deterioration in cardiac parameters such as stroke volume and cardiac output. These findings suggest a dynamic and possibly age-dependent role for miR-378a in modulating cardiac performance. Interestingly, despite evidence of impaired contractility and adverse remodeling in younger animals, some parameters appear normalized or compensated in older mice, hinting at either an adaptive response or a shift in the underlying mechanisms of dysfunction over time. In line with such observation, in our previous study we noticed that the effect miR-378a knockout in murine model of Duchenne muscular dystrophy (*mdx* strain) is less pronounced in 6 month old mice in comparison to their younger, 3 month old, counterparts [22].

Although our transcriptomic and proteomic analyses in hiPSC-CM and murine hearts did not identify many individual molecules with statistically significant changes, integration of these datasets uncovered substantial alterations in biological pathways. GO and KEGG pathway analyses pointed toward disrupted regulation of muscle contraction, post-transcriptional gene expression, and notably, energy metabolism. The modest overlap between the transcriptome and proteome datasets is not unexpected, given the known challenges in correlating mRNA and protein levels, particularly in the context of miRNA-mediated regulation which often involves translational repression rather than mRNA degradation. Of note, several possible targets of miR-378a such as YY1 were upregulated which opens new avenues for deeper investigation of the miR-378a-mediated pathways in the heart. Interestingly, transgenic cardiac overexpression of YY1 was reported to induce pathological hypertrophy, but only in male mice [23]. In accordance with this observation, YY1 level was also higher in male but not female heart failure patients in comparison to control counterparts [23]. Taking into consideration that we used in our study only male mice and two out of three hiPSC lines were male donor-derived, upregulation of YY1 might be another mechanism by which miR-378a regulates hypertrophic growth of cardiomyocytes. So far, IGF1R-MAPK pathway was associated with miR-378a-mediated effect on this process [7,8,10]. For instance, we previously reported upregulation of pAKT as well as total and pERK in miR-378a-KO hiPSC-CM which was followed by increased level and nuclear localization of NFATc3 [8]. In line with these observations in the current study IGF1R receptor, pAKT and pERK were more abundant in miR-378a-/- mice already at 6 weeks (Fig. 1C). Whether YY1 is another factor directly targeted by miR-378a and involved in miR-378a-mediated regulation of cardiac hypertrophy is still to be confirmed but our combined analysis of mouse and human omics dataset may provide robust source of more of such putative factors.

A major finding of this study, reflected by the combined omics, is the miR-378a-mediated regulation of metabolic processes in human cardiomyocytes. First, Seahorse analysis revealed broad suppression of mitochondrial function, including basal respiration, ATP production, and spare respiratory capacity, consistent with bioenergetic insufficiency. These functional deficits were accompanied by morphological abnormalities in mitochondria and reduced mitochondrial content. Mechanistically, our data suggest that miR-378a is required for a balanced mitophagy–biogenesis axis. Elevated levels of mitophagy initiators (e.g., Parkin and p62) and impaired formation of autolysosomes upon CCCP stimulation point toward defective mitophagic flux. Simultaneously, markers of mitochondrial biogenesis such as TFAM were suppressed, and restoration of miR-378a expression partially reversed these effects. Notably, while PGC1α protein levels remained unchanged, the decrease in TFAM and mitochondrial DNA content suggests that downstream biogenic signaling is nonetheless compromised. Together, these findings identify miR-378a as an important coordinator of mitochondrial turnover in human cardiomyocytes – a novel and physiologically significant role with implications for cardiac energy homeostasis and cardiomyocyte maturation. The latter process is not limited solely to organ growth but primarily to numerous changes, both genetic and metabolic. Cardiac metabolism during fetal development relies primarily on glycolysis, which is associated with reduced oxygen conditions and a lower demand for efficient blood pumping compared to postnatal period. The heart matures during the first decade of life, adapting its metabolism by switching from glycolysis to β-oxidation of fatty acids, as well as changing the isoforms of proteins that build sarcomeres [24]. Interestingly, Kuppusamy et al. observed that long term culture as well as formation of engineered heart tissues which both induced substantial maturation of hESC-CM was associated with upregulation of several miRNAs and concomitant downregulation of their potential targets [6]. Among these molecules let-7 and miR-378a were one of the most profoundly elevated indicating their important role in regulating hESC-CM phenotype [6]. Further experiments revealed that let7 promoted maturation of human cardiomyocytes by modulating their metabolic state. Thus, increase of miR-378a expression with time and our current results demonstrating its effect on hiPSC-CM metabolism may indicate that this miRNA is also involved in the maturation of human cardiomyocytes.

In addition to its effects on mitochondrial respiration, the knockout of miR-378a altered glucose handling in cardiomyocytes. Despite increased glucose uptake – presumably as a compensatory response – glycogen accumulation and the activity of glycolytic enzymes such as hexokinase and LDH were decreased. These seemingly paradoxical results are indicative of impaired glucose handling, where increased substrate availability is not efficiently utilized or stored. Upregulation of GLUT1 and AMPKγ supports a stress-induced metabolic adaptation, but elevated GSK3α/β levels may inhibit glycogen synthesis, as supported by our PAS staining results. This regulatory profile mirrors previous observations of miR-378a activity in non-cardiac tissues, where it modulates both mitochondrial and glycolytic energy pathways [2,18,25]. Our data extend these findings to human cardiomyocytes and underscore the metabolic vulnerability imposed by miR-378a loss.

Another important observation from this study is the reduced angiogenic capacity of miR-378a-KO cardiomyocytes. Despite higher VEGF secretion, conditioned media from these cells significantly impaired endothelial tube formation in both 2D assay. This discrepancy suggests the presence of anti-angiogenic factors or altered VEGF bioavailability/sequestration in the miR-378aKO secretome. The complexity of miRNA-regulated secretory profiles could explain such outcomes and warrants further investigation. The reduced angiogenic capacity aligns with our earlier findings in skeletal muscle and diabetic heart models [4,10], suggesting a consistent role for miR-378a in modulating angiogenesis across tissues.

Our study aimed also to evaluate whether overexpression of miR-378a in hiPSC-CM could enhance their therapeutic potential in the context of myocardial infarction. Although prior observations highlighted the role of miR-378a in supporting metabolic homeostasis and angiogenic competence of hiPSC-CM, our *in vivo* findings demonstrate that while transplantation of both control and miR-378a-overexpressing cardiomyocytes significantly improved cardiac function at the 4-week time point, miR-378a overexpression did not provide an additive benefit. Significant improvement in LV systolic function, as indicated by increased ejection fraction and fractional shortening, was observed in both cell-treated groups, accompanied by a reduction in left ventricular end-diastolic and end-systolic diameters, highlighting that hiPSC-CM delivery not only enhances contractile performance but also attenuates pathological ventricular remodeling after myocardial infarction. However, the lack of an additional effect in the miR-378a-overexpressing group suggests that either the baseline benefits of hiPSC-CM therapy reach a ceiling effect or that miR-378a’s pro-regenerative influence is context-dependent and insufficient on its own to further enhance cardiac repair in this setting.

Of note, the level of miR-378a overexpression in hiPSC-CM after transduction with AAV6-miR-378a vectors was not very high which may further limit its possible therapeutic effect. On the other hand, in our previous study evaluating the effect of heme oxygenase-1 (HO-1) and SDF-1 upregulation on the therapeutic potential of hiPSC-CM in the same animal model of MI we also did not observe the additive effect of introduced factors [11], thus we cannot exclude that in the applied experimental settings the window for further improvement of heart function is highly limited.

### 4.1. Limitations and further perspective

Despite these comprehensive findings, several limitations must be acknowledged. First, proteomic analyses yielded relatively few statistically significant hits in isolation, reflecting the challenges posed by inter-donor variability in hiPSC lines and the sensitivity of mass spectrometry-based detection. Second, while we used multiple hiPSC lines to minimize donor-related artifacts, gene editing and clonal selection may still introduce subtle confounders. Finally, the metabolic assays were limited to in vitro models; validating these results *in vivo* will be an important next step. Further research should also focus on identifying direct targets of miR-378a that mediate its metabolic and angiogenic effects. Moreover, therapeutic modulation of miR-378a may hold promise in conditions characterized by energetic deficiency or impaired vascularization, such as chronic heart failure.

## Authors’ contributions

A.M., I.W., K.P., J.K., M.T., N.L., U.F-S. and J.S. performed experiments. L.S. and T.B. performed the proteome analysis. O.W. prepared the samples for TEM imaging. E.P., G.M. T.G. and G.Y. analysed the transcriptome and proteome data. I.K. and K.S. assisted with AAV vector production. M.K. and E.P. helped with data interpretation. J.S, A.M., U.F-S,., and J.D. designed the experiments. A.M., U.F-S. and J.S. prepared figures and wrote the manuscript. J.D. reviewed the manuscript. J.D., U.F-S and J.S. supervised the work. All authors reviewed and accepted the manuscript.

## Acknowledgments

The authors would like to acknowledge Agnieszka Andrychowicz-Róg^†^ and Joanna Uchto-Bajołek from the Department of Medical Biotechnology (Jagiellonian University in Kraków, Poland) for administrative assistance.

## Availability of data and materials

The transcriptomic and proteomic data included in this article will be available in the Dryad Digital Repository upon acceptance.

## Declarations

### Ethics approval and consent to participate

This study was approved by the following ethical committees: *Title:* Investigation of the effect of microRNA-378 on the pathophysiology of diabetic cardiomyopathy in a murine model; *Name of the institutional approval committee:* The Second Local Ethical Committee for Animal Research in Krakow, Poland; *Approval number:* 108/2021; *Date of approval:* 08.04.2021. *Title:* Regulation of microRNA-378 as a novel strategy to enhance the maturity and therapeutic potential of human cardiomyocytes derived from induced pluripotent stem cells; *Name of the institutional approval committee:* The Second Local Ethical Committee for Animal Research in Krakow, Poland; *Approval number:* 358/2021; *Date of approval:* 08.12.2021. *Title:* Investigation of the effect of microRNA-378a deficiency on cardiac vascularization in mice; *Name of the institutional approval committee:* The Second Local Ethical Committee for Animal Research in Krakow, Poland; *Approval number:* 354/2022; *Date of approval:* 1.12.2022. *Title:* Generation of human induced pluripotent stem cells from peripheral blood mononuclear cells, followed by their differentiation into cardiomyocytes, myoblasts, and endothelial cells; *Name of the institutional approval committee:* Jagiellonian University Bioethical Committee in Krakow, Poland; *Approval number:* 122.6120.303.2016; *Date of approval:* 24.11.2016.

### Informed consent

Informed consent was obtained from all human participants prior to their inclusion in the study. All participants provided written consent, confirming their willingness to participate and their understanding of the study.

### Consent for publications

All authors confirm their consent for publication.

### Declaration of generative AI and AI-assisted technologies in the writing process

During the preparation of this manuscript, ChatGPT was used solely for language editing and improving readability. The authors reviewed and edited the content and take full responsibility for the final version of the manuscript.

