## Supplementary Material for "miR-378a Controls Cardiomyocyte Metabolism and Angiogenic Signaling"

### Supplementary Figures

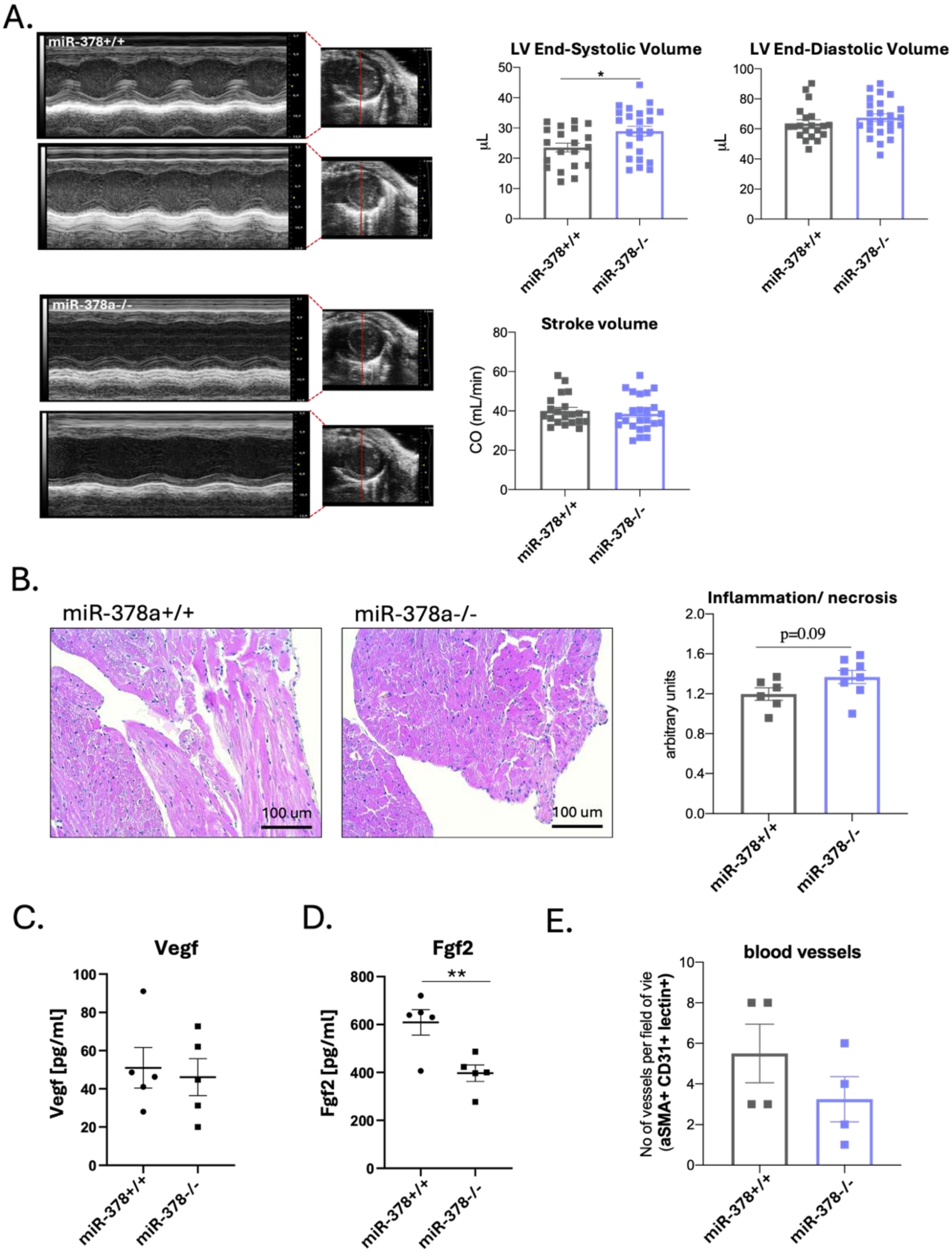


***Supplementary Figure 1. A.*** *Representative echocardiographic tracings recorded in miR-378a+/+ (upper left) and miR-378a-/- (bottom left) mice. Right panel: LV End-Systolic Volume, LV End-Diastolic Volume and Stroke Volume (CO – cardiac output) in 10 – 13-week-old miR-378a+/+ and miR-378a-/- mice. n=20 miR-378a+/+ mice and n=24 miR-378-/- mice.* ***B.*** *Representative cardiac sections from 11 – 12-week-old miR-378+/+ and miR-378a-/- mice (left panel). The level of inflammation/necrosis was calculated based on these sections (right panel). Scale bar = 100 μm.* ***C – D.*** *ELISA-based analysis of Vegf (**C.**) and Fgf2 (**D.**) protein level in miR-378a+/+ and miR-378a-/- heart lysates isolated from 11 – 12-week-old mice. n=5.* ***E.*** *Quantification of αSMA-, CD31- and lectin-positive vessels in cardiac sections prepared from the miR-378a+/+ and miR-378a-/- hearts of 11 – 12-week-old mice, n=4. *p<0.05, **p<0.01, unpaired two-tailed Student's t-test.*

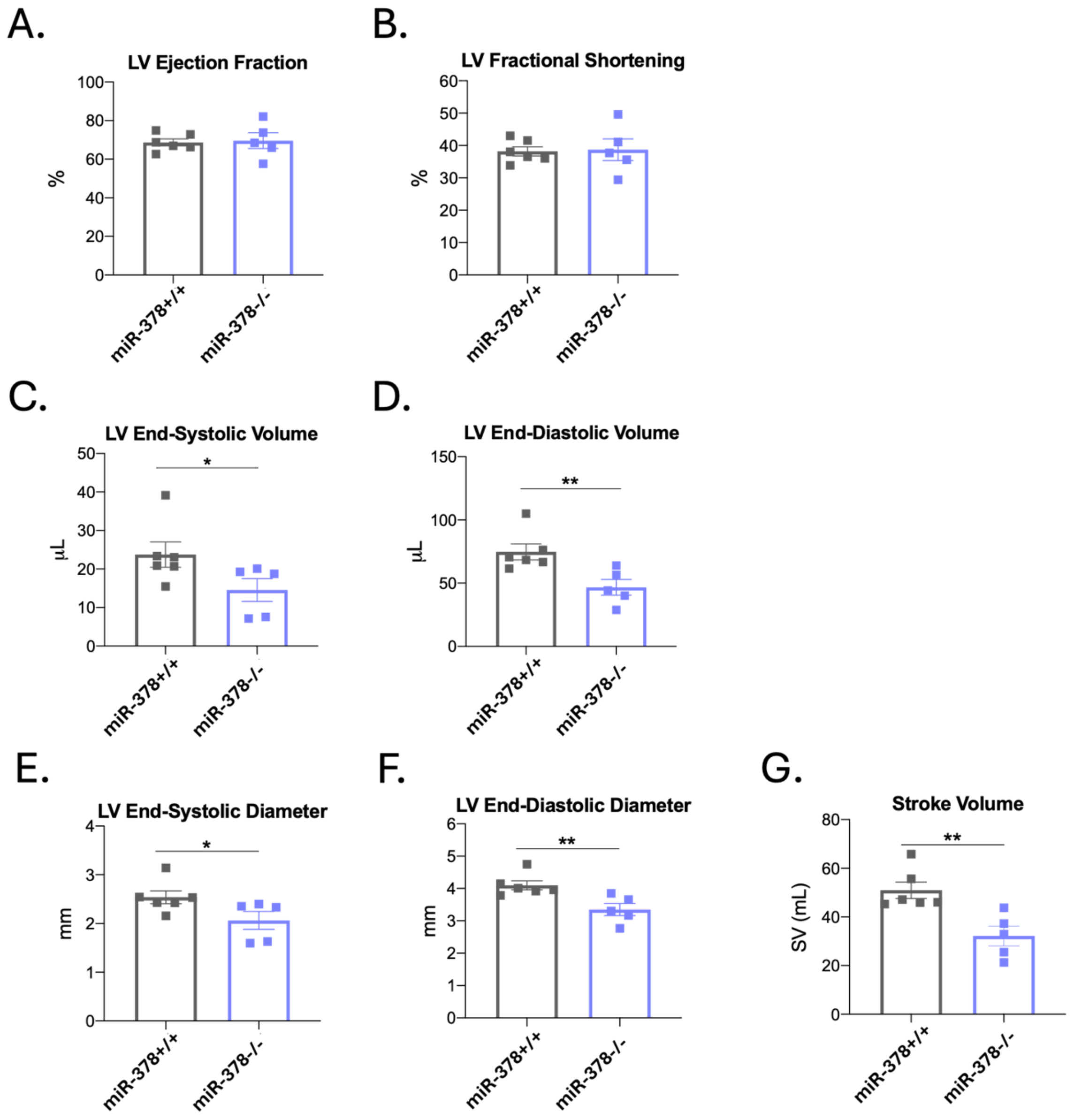


***Supplementary Figure 2. A – H.*** *LV Ejection Fraction (EF) (**A.**), LV Fractional Shortening (FS) (**B.**), LV End-Systolic Volume (**C.**), LV End-Diastolic Volume (**D.**), LV End-Systolic Diameter (**E.**), LV End-Diastolic Diameter (**F.**) and Stroke Volume (SV) (**G.**) in 17 month old miR-378a+/+ and miR-378a-/- mice. n=5-6, *p<0.05, **p<0.01, unpaired two-tailed Student's t-test or Mann–Whitney U test.*

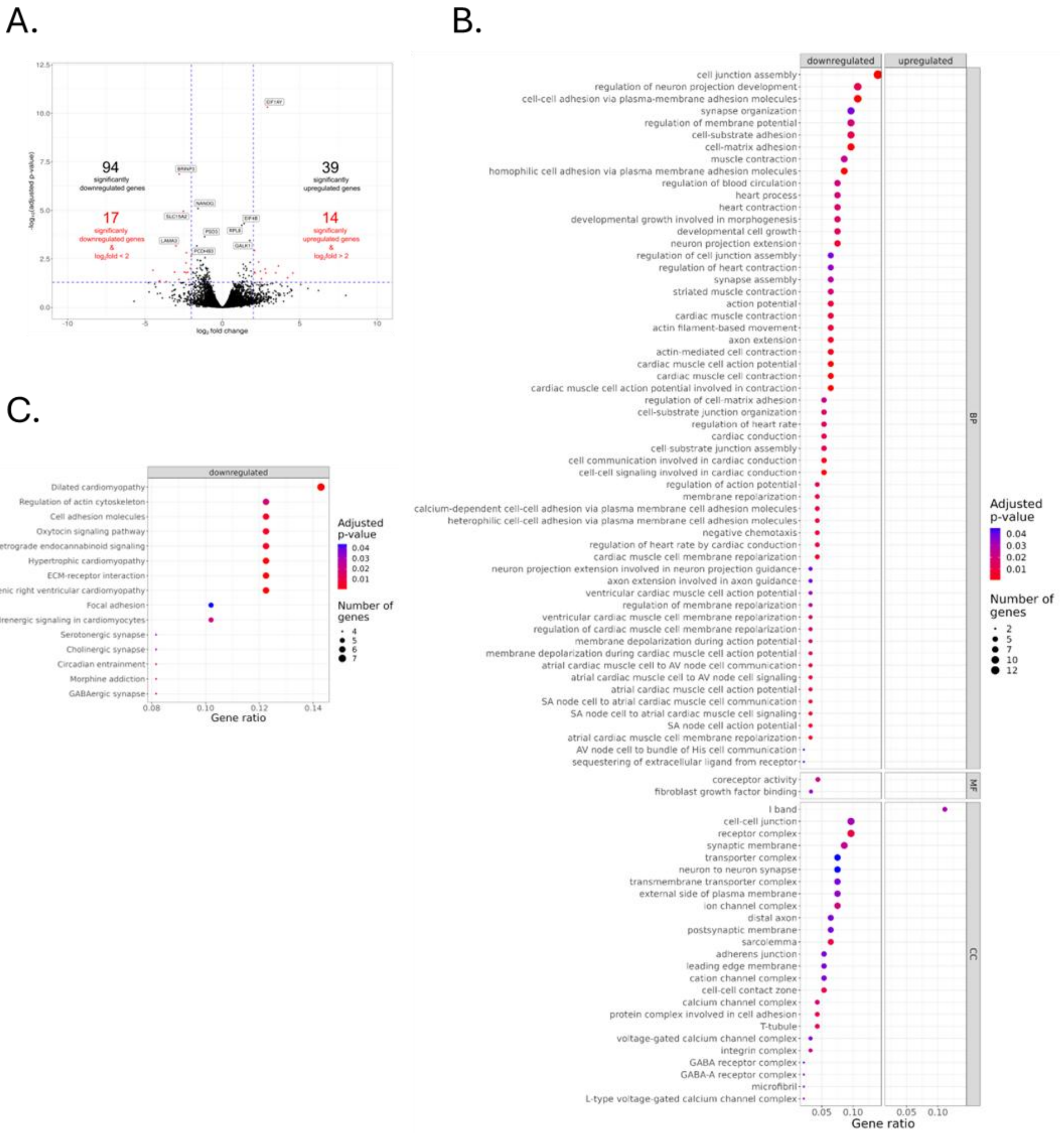


***Supplementary Figure 3. A.*** *Volcano plot of the transcriptomic differential expression analysis of miR-378aKO hiPSC-CM in comparison to their control counterparts. Genes with* $|log_2FoldChange| > 2$ *and adjusted p-value < 0.05 are shown in red. The names of top 10 significant genes are labelled. Blue dotted lines represent* $|log_2FoldChange| = 2$ *(vertical) and the adjusted p-value of 0.05 (horizontal).* ***B.*** *Dot plot of GO-terms over-representation analysis of transcriptomic dataset from control and miR-378aKO hiPSC1-CM. Only genes with significantly altered expression (adjusted p-value < 0.05) were included. BP – biological processes, MF – molecular function, CC – cellular components.* ***C.*** *Dot plot of KEGG over-representation analysis of control and miR-378aKO hiPSC1-CM transcriptomes. Only genes with significantly altered expression (adjusted p-value < 0.05) were included. No significantly over-represented KEGG pathways were detected for upregulated genes.*

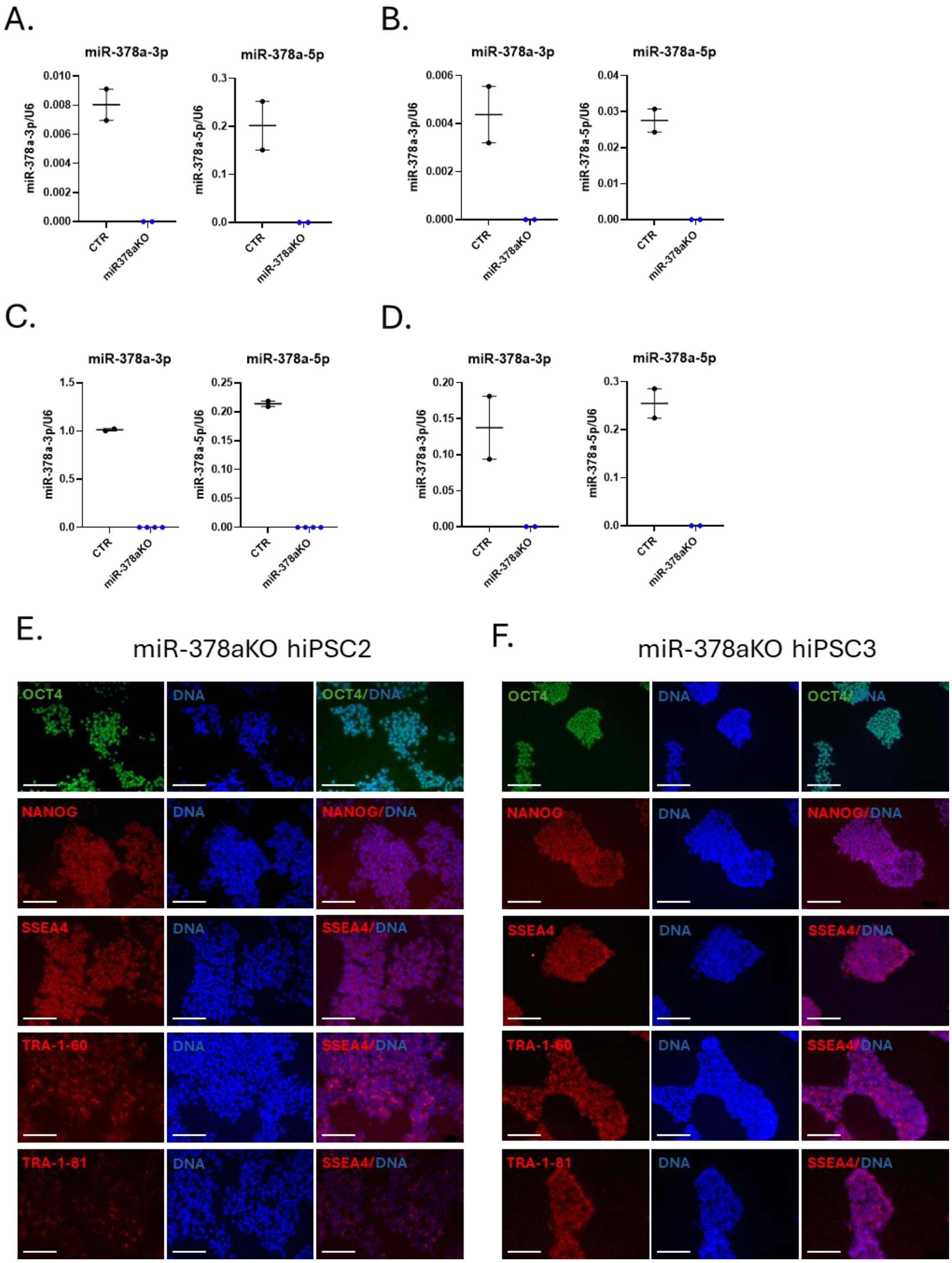


***Supplementary Figure 4. A – D.*** *qPCR analysis of miR-378a-3p and miR-378a-5p expression in control and miR-378aKO hiPSC2 (**A.**) and hiPSC3 (**B.**) as well as control and miR-378aKO hiPSC2-CM (**C.**) and hiPSC3-CM (**D.**). **F – G.** Pluripotency marker expression in miR-378aKO hiPSC2 (**F.**) and miR-378aKO hiPSC3 (**G.**): OCT4 (green), NANOG (red), SSEA4 (red) and TRA-1-60 (red) and TRA-1-81 (red). Hoechst33342 (blue) was used to stain nuclei. Scale bar = 10 µm.*

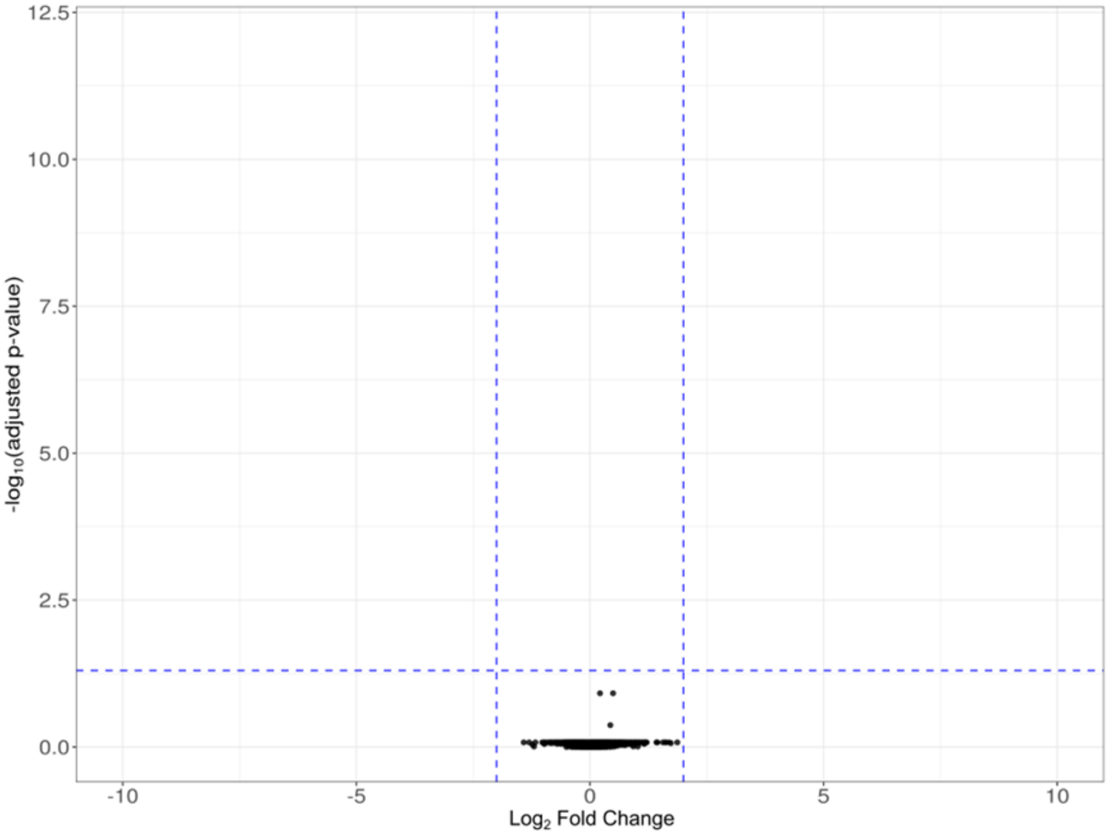


***Supplementary Figure 5.*** *Vulcano plot presenting the proteomic differential expression analysis in miR-378aKO hiPSC-CM vs their respective control counterparts. Blue dotted lines represent |$log_2$FoldChange| = 2 (vertical) and the adjusted p-value of 0.05 (horizontal).*

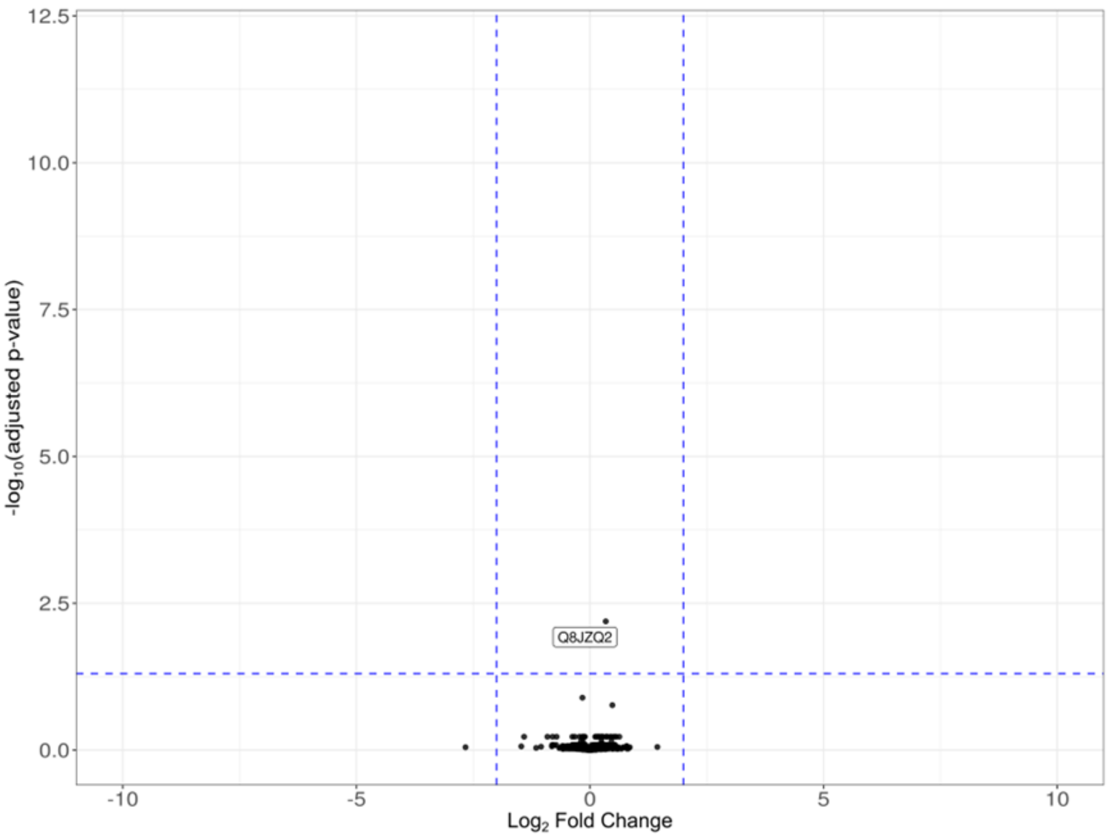


***Supplementary Figure 6.*** *Vulcano plot presenting the proteomic differential expression analysis in miR-378a-/- murine hearts vs their respective control counterparts. Blue dotted lines represent |log$_2$FoldChange| = 2 (vertical) and the adjusted p-value of 0.05 (horizontal).*

| DIRECTION | GENE_HUMAN | GENE_NAME | PROTEIN_HUMAN | PROTEIN_MOUSE |
|---|---|---|---|---|
| DOWN | CHRM2 | cholinergic receptor muscarinic 2 | P08172 | Q9ERZ4 |
| DOWN | DBT | dihydrolipoamide branched chain transacylase E2 | P11182 | P53395 |
| DOWN | PYGB | glycogen phosphorylase B | P11216 | Q8CI94 |
| DOWN | TPR | translocated promoter region, nuclear basket protein | P12270 | F6ZDS4 |
| DOWN | IVD | isovaleryl-CoA dehydrogenase | P26440 | Q9JHI5 |
| DOWN | NDUFV3 | NADH:ubiquinone oxidoreductase subunit V3 | P56181 | Q8BK30 |
| DOWN | SORD | sorbitol dehydrogenase | Q00796 | Q64442 |
| DOWN | AARS2 | alanyl-tRNA synthetase 2, mitochondrial | Q5JTZ9 | Q14CH7 |
| DOWN | CPSF7 | cleavage and polyadenylation specific factor 7 | Q8N684 | Q8BTV2 |
| DOWN | FYCO1 | FYVE and coiled-coil domain autophagy adaptor 1 | Q9BQS8 | Q8VDC1 |
| DOWN | OPA3 | outer mitochondrial membrane lipid metabolism regulator OPA3 | Q9H6K4 | Q505D7 |
| DOWN | GDE1 | glycerophosphodiester phosphodiesterase 1 | Q9NZC3 | Q9JL56 |
| DOWN | CAB39 | calcium binding protein 39 | Q9Y376 | Q06138 |
| UP | ESYT2 | extended synaptotagmin 2 | A0FGR8 | Q3TZZ7 |
| UP | EIF3D | eukaryotic translation initiation factor 3 subunit D | O15371 | O70194 |
| UP | PSMD3 | proteasome 26S subunit, non-ATPase 3 | O43242 | P14685 |
| UP | MYO1B | myosin IB | O43795 | P46735 |
| UP | SNX2 | sorting nexin 2 | O60749 | Q9CWK8 |
| UP | RTN2 | reticulon 2 | O75298 | O70622 |
| UP | SEC31A | SEC31 homolog A, COPII coat complex component | O94979 | Q3UPL0 |
| UP | OXSR1 | oxidative stress responsive kinase 1 | O95747 | Q6P9R2 |
| UP | SOD1 | superoxide dismutase 1 | P00441 | P08228 |
| UP | HSP90AA1 | heat shock protein 90 alpha family class A member 1 | P07900 | P07901 |
| UP | EEF1B2 | eukaryotic translation elongation factor 1 beta 2 | P24534 | O70251 |
| UP | YY1 | YY1 transcription factor | P25490 | Q00899 |
| UP | AKT1 | AKT serine/threonine kinase 1 | P31749 | P31750 |
| UP | BAG6 | BAG cochaperone 6 | P46379 | Q9Z1R2 |
| UP | SERPINH1 | serpin family H member 1 | P50454 | P19324 |
| UP | RPS7 | ribosomal protein S7 | P62081 | P62082 |
| UP | GRB2 | growth factor receptor bound protein 2 | P62993 | Q60631 |
| UP | LMNB2 | lamin B2 | Q03252 | P21619 |
| UP | CSTF3 | cleavage stimulation factor subunit 3 | Q12996 | Q99LI7 |
| UP | EIF4H | eukaryotic translation initiation factor 4H | Q15056 | Q9WUK2 |
| UP | CMBL | carboxymethylenebutenolidase homolog | Q96DG6 | Q8R1G2 |
| UP | PRMT1 | protein arginine methyltransferase 1 | Q99873 | Q9JIF0 |

***Supplementary Figure 7.*** *List of downregulated (blue) and upregulated (red background) putative miR-378a targets in miR-378a-deficient murine hearts and hiPSC-CM found after combined analysis of all omic datasets and miRTarBase database.*

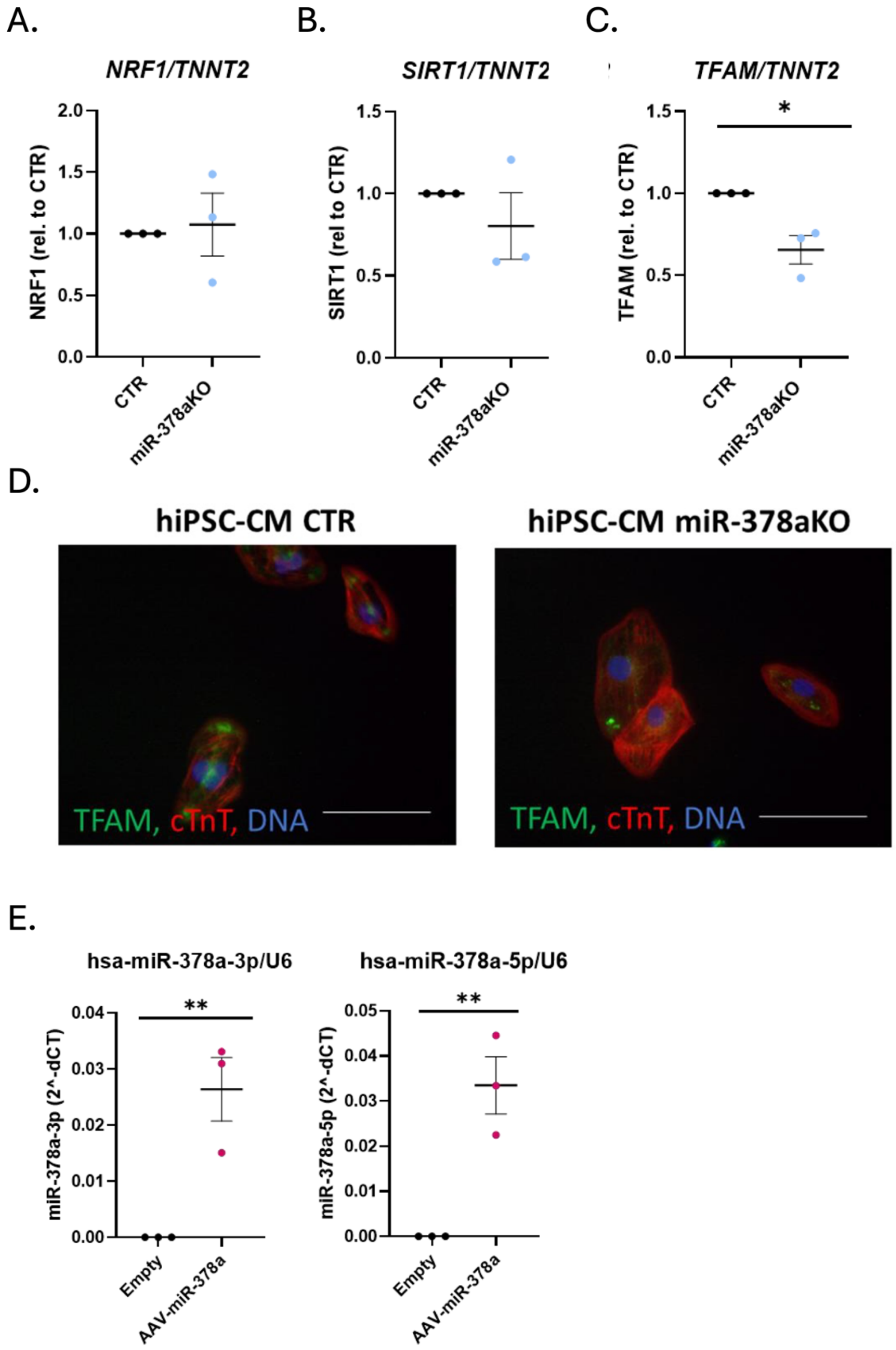


***Supplementary Figure 8. A – C** qPCR analysis of NRF1 (**A.**), SIRT1 (**B.**) and TFAM (**C.**) in control (CTR) and miR-378aKO hiPSC2-CM. TNNT2 was used as a reference control, n=3, *p<0.05, unpaired two-tailed Student's t-test. **D.** Representative images from immunofluorescent analysis of TFAM (green) and cardiac troponin T (cTnT) expression in control (CTR) and miR-378aKO hiPSC1-CM. Nuclei (DNA) were stained in blue. Scale bar = 10 µm. **E.** qPCR analysis of miR-378a-3p (left panel) and miR-378a-5p (right panel) expression in miR-378aKO hiPSC-CM transduced with AAV6-empty (Empty) and AAV6-miR-378a (AAV-miR-378a) vectors. U6 was used as a reference control. n=3, **p<0.01, unpaired two-tailed Student's t-test.*

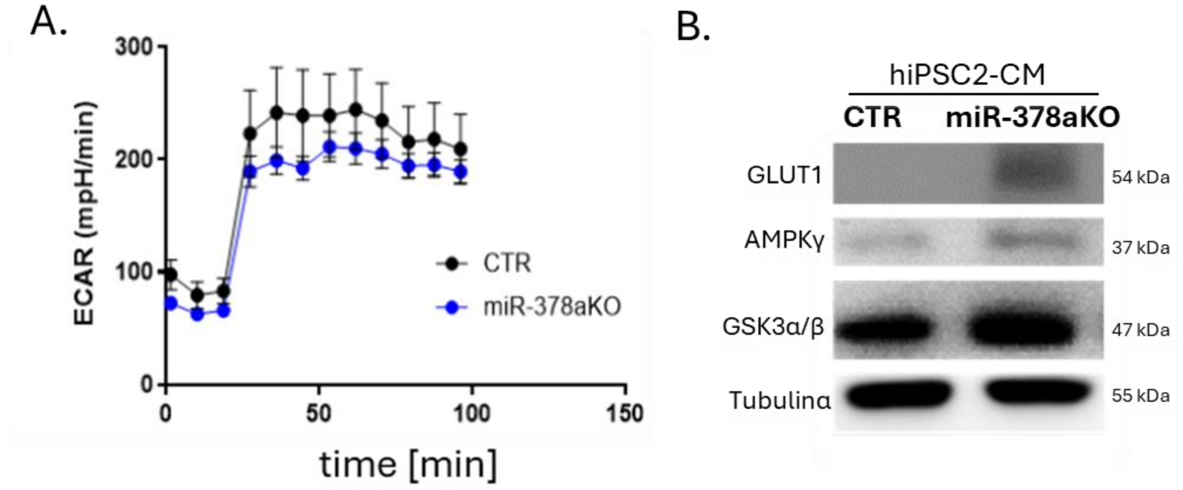


***Supplementary Figure 9. A.*** *Extracellular acidification rate (ECAR) during Seahorse XF Mito Stress assay in control (CTR) and miR-378aKO hiPSC1-CM.* ***B.*** *Western blot analysis of GLUT1 and GSK3α/β in control (CTR) and miR-378aKO hiPSC2-CM. α Tubulin was used as a protein loading control.*

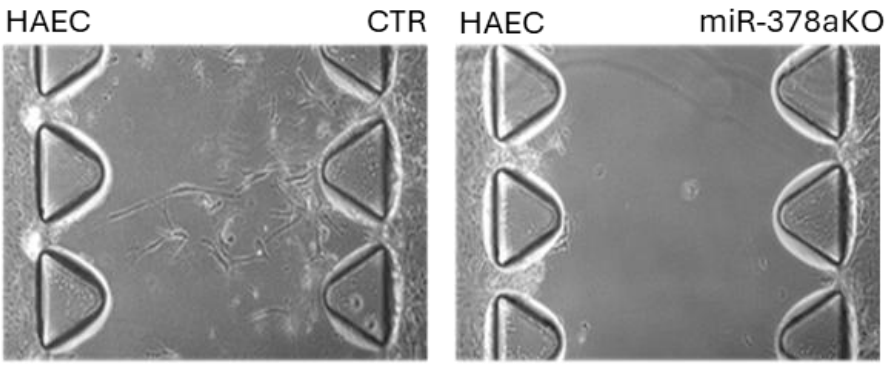


***Supplementary Figure 10.*** *Representative images of idenTx 3 chips with HAEC seeded in the left lateral channel and either control (CTR) or miR-378aKO hiPSC-CM in the right lateral channel, n=2.*

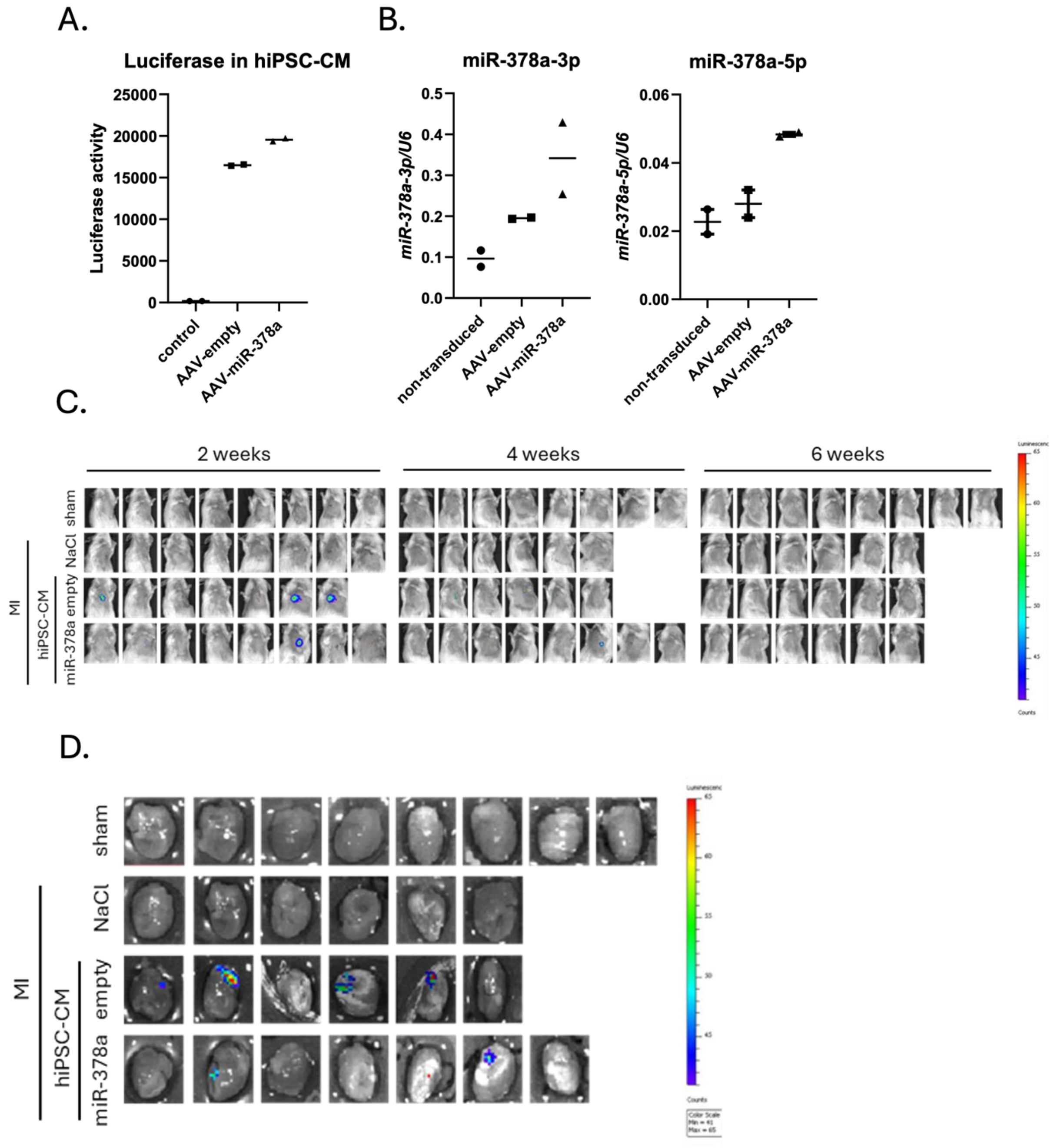


***Supplementary Figure 11. A.*** *Luciferase activity in hiPSC1-CM transduced with AAV6-Luc and either AAV6-empty or AAV6-miR-378a. Non-transduced hiPSC1-CM served as a control.* ***B.*** *qPCR analysis of miR-378a-3p (left panel) and miR-378a-5p (right panel) in hiPSC1-CM transduced with AAV6-Luc and either AAV6-empty or AAV6-miR-378a. Non-transduced hiPSC1-CM served as a control.* ***C.*** *Bioluminescent signal collected intravitally from the chest of all animals used in the experiment two, four and six weeks after MI induction and cell administration.* ***D.*** *Bioluminescent signal collected from isolated hearts six weeks after MI induction and cell administration. empty – hiPSC1-CM transduced with AAV6-empty vectors; miR-378a – hiPSC1-CM transduced with AAV6-miR-378a vectors.*

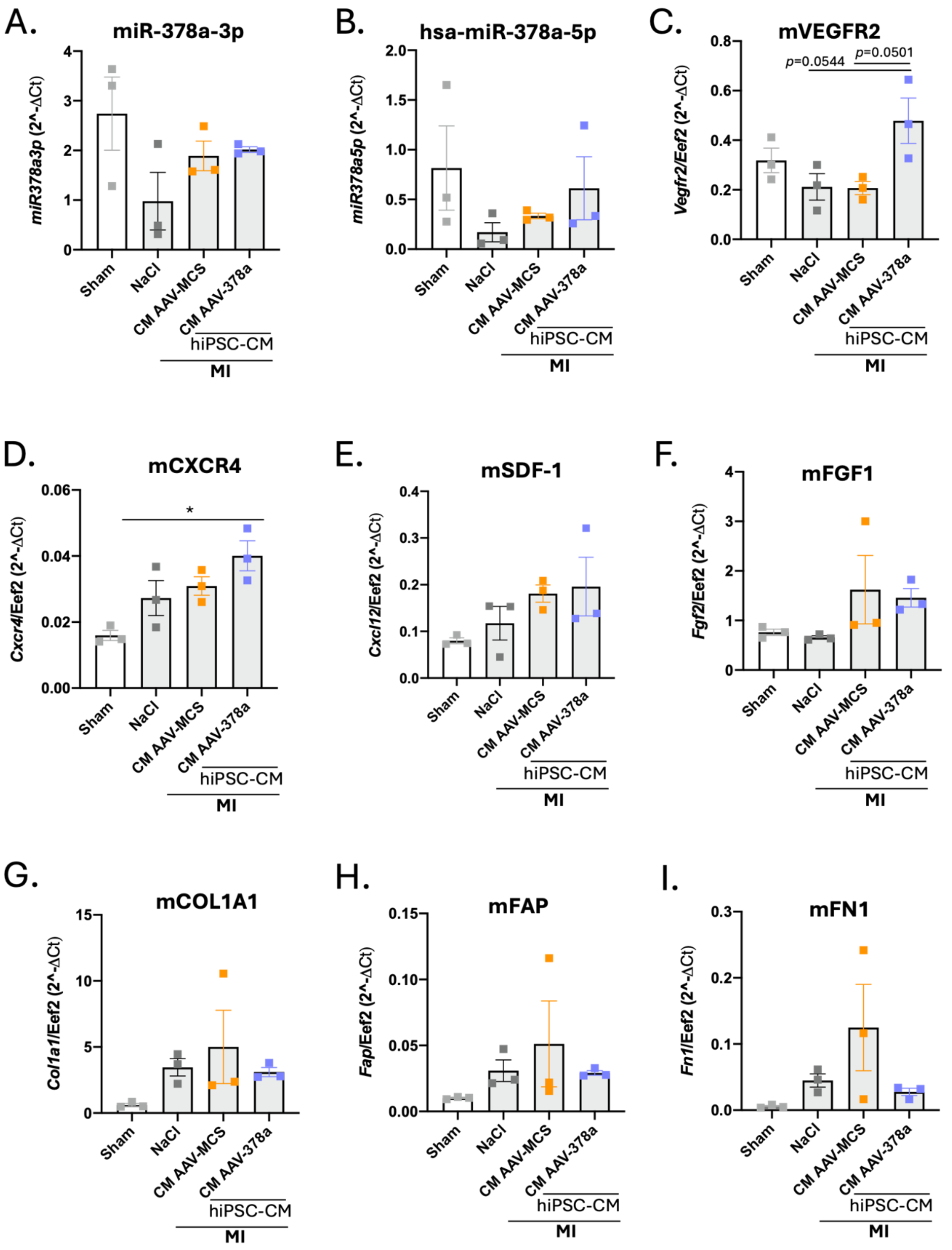


***Supplementary Figure 12. A – J.*** *qPCR analysis of miR-378a-3p (**A.**), miR-378a-5p (**B.**) as well as murine (m) Kdr (encoding Vegfr2) (**C.**), Cxcr4 (**D.**), Cxcl12 (**E.**), Fgf1 (**F.**), Col1a1 (**G.**), Fap (**H.**) and Fn1 (**I.**) expression in the heart lysates isolated from mice subjected to sham operation (sham) as well as MI induction followed by either vehicle (NaCl) or hiPSC1-CM transduced with AAV6-empty vectors (AAV-empty) or hiPSC1-CM transduced with AAV6-miR-378a vectors (AAV-miR-378a) administration. n=3, *p<0.05 vs sham group, one-way ANOVA, followed by Tukey's post hoc test.*

**Supplementary Methods**

### 1.1. CRISPR/Cas9 gene editing

Line 1 hiPSC with miR-378a knockout (miR-378aKO) was previously generated by us [1] using CRISPR/Cas9 gene editing. The same method was used to derive hiPSC miR378aKO cells from hiPSC line 2 and 3 with a different set of sgRNAs (sgRNA1: 5'-GATCTTTGCCGGCCCAACTT-3'; sgRNA2: 5'-AGTTCAAATGGCTTGCTCCG-3') to address the issue of possible off-target effects of the applied system. Briefly, 500,000 cells were nucleofected with the sgRNA1-introduced pSpCas9-2A-Puro plasmid (Addgene plasmid #62988 [2]) and the sgRNA2-introduced lentiCRISPR v2-Blast plasmid (Addgene plasmid #83480). Nucleofection was performed using the Human Stem Cell Nucleofector Kit 1 (Lonza) according to the manufacturer's instructions. 24 hours after nucleofection, antibiotic selection was performed to eliminate cells that did not uptake the plasmids. Two antibiotics were used for selection: puromycin (Sigma-Aldrich) (0.7 μg/mL for hiPSC2 and 0.3 μg/mL for hiPSC3) and blasticidin (InvivoGene) (10 μg/mL for hiPSC2 and 20 μg/mL for hiPSC3). After 24 hours, the medium was changed to E8 without antibiotics. In the next step, 500 (hiPSC2) or 100 (hiPSC3) cells were seeded on a 10 cm diameter dish to allow the formation of colonies derived from single cells and cultured for 10–14 days. The colonies obtained in this way were collected, subjected to genetic screening as described previously [1] and those with confirmed deletion of *MIR378A* locus were expanded and used in further experiments.

### 1.2. Cardiac differentiation

Cardiac differentiation of hiPSC was performed according to the protocol previously applied in our studies[1,3–5]
and described by Lian et al.[6]. Briefly, 30,000 cells were seeded per well of a 24-well plate and cultured until 90 – 100% confluency. Thereafter, the medium was changed to RPMI 1640 (Sigma Aldrich) supplemented with B27 without insulin (ThermoFisher Scientific) (RPMI/B27-Ins) and hiPSC were stimulated with an optimized concentration (Table 1) of a small molecule GSK3α/β kinase inhibitor – CHIR 99021 (Sigma Aldrich). After 24 hours, the medium was changed to RPMI/B27-Ins, and 72 h after the first stimulation, to induce differentiation towards cardiac mesoderm, an optimized concentration (Table 1) of a small molecule WNT pathway inhibitor – IWR-1 (Sigma Aldrich) was used in the conditioned medium (1:1 mixture of fresh RPMI/B27-Ins and the medium from the cells). After 48 hours, the medium was changed to RPMI/B27-Ins. From day 7 of differentiation, cells were cultured in RPMI 1640 medium with B27 supplement (ThermoFisher Scientific) (RPMI/B27), refreshed

every 3 days. Between day 13 and 19, metabolic selection was introduced using RPMI 1640 without glucose (RPMI 1640 no glucose, ThermoFisher Scientific) supplemented with B27 and 4 mM sodium lactate (Sigma-Aldrich). Subsequently, cells were collected using Multi Tissue Dissociation reagent 3, (Miltenyi Biotec) according to the manufacturer's protocol. After seeding on Geltrex$^{TM}$-coated wells, culture was continued in RPMI/B27 medium, and experiments were performed between day 25 and 30 of differentiation.

*Table 1. Optimized concentrations of CHIR99021 and IWR-1 used for cardiac differentiation of hiPSC.*

| Line | Genotype | CHIR90221 | IWR-1 |
|---|---|---|---|
| **hiPSC 1** | control | 12 µM | 12 µM |
| | miR-378aKO | | |
| **hiPSC 2** | control | 8 µM | 5 µM |
| | miR-378aKO | | |
| **hiPSC 3** | control | 8 µM | 5 µM |
| | miR-378aKO | | |

#### 1.3. Production of AAV vectors

Production of AAV6-CMV-miR-378a, AAV6-CMV-Luc and AAV6-CMV-empty vectors was performed as previously described[7]. Briefly, AAV-293 cells were transfected with either pAAV-CMV-miR-378a (kindly gifted by Prof. Mauro Giacca, International Centre for Genetic Engineering and Biotechnology, Trieste, Italy), pAAV-CMV-empty or pAAV-CMV-Luc together with helper pDGM6 plasmid (Addgene plasmid #110660[8]) using PEI MAX (Polysciences) transfection reagent. 72 hours after transfection, vector isolation was performed using freezing/thawing cell lysis followed by the incubation with HS nuclease (Mi Bio Tech) and ultracentrifugation in an iodixanol gradient (Sigma-Aldrich).

Quantitative PCR was used to determine virus titer. In the first step, 5 µL of purified vectors were digested with DNase I (ThermoFisher Scientific). For this purpose, a mixture consisting of 5 µL of vectors, 10 µL of 10x concentrated DNaseI buffer (ThermoFisher Scientific), 1 µL of DNase I (1 U/µl) and 84 µl of water was prepared and incubated for 1.5 h at 37°C. DNase I was then inactivated by adding 25 mM EDTA and incubating for 10 min at 65°C. In the next step, proteinase K (A&A Biotechnology) digestion was performed and incubation with shaking for 1 h at 37°C. Next, DNA extraction and purification from vectors was carried out. 500 µl of phenol-chloroform-isoamyl alcohol solution mixed in a ratio of 25:24:1 (Sigma-Aldrich) was added and mixed vigorously for 1 min and then centrifuged (5 min, 12,000 × g). After centrifugation, the upper aqueous phase was collected and transferred to a fresh tube, then 10 µL of 5 M NaCl solution and 1 mL of 100% ethanol (POCH) were added, vortexed and incubated overnight at -20°C. The next day, it was centrifuged (15 min, 14,000 × g) and purified using 70% ethanol. Standard curve of 10-fold dilutions (from 10 ng/µl to $10^{-5}$ ng/µl) of the linearized fragment of the pAAV-CMV-empty plasmid and 10-, 100- and 1000-fold dilutions of the isolated DNA were prepared. A mixture of 10 µl of TaqMan Universal Master Mix II (Applied Biosystem), 1 µl of 5 µM Forward primer (5'-GTGGGAGGTCTATATAAGC-3'), 1 µl of 5 µM Reverse primer (5'-GACCTCCATAGAAGACAC-3'), 1 µl of 2 µM probe (5' 6-FAM-AGTGAACCGTCAGATCGCCT-BHQ-1 3') and 2 µl of DNA was prepared. The reaction was performed using StepOnePlus TM Real-time PCR Systems thermal cycler (Applied Biosystems) with the following parameters: initial denaturation and enzyme activation (95°C, 10 min), followed by a denaturation cycle repeated 40 times (95°C, 15 s) and primer annealing/extension/fluorescence detection (60°C, 1 min). Viral titer was calculated based on the standard curve. hiPSC-CM were transduced with AAV6-Luc vectors at the multiplicity of infection (MOI) = 4000 while with AAV6-empty and AAV6-miR-378a at MOI=10000 virus particles/cell.

#### 1.4. Endothelial cell (EC) tube formation assay

Tube formation assay was performed according to the previously described protocol[9]. Briefly, Human Aortic Endothelial Cells (HAEC) were seeded on 96-well plates pre-coated with 50 µL of growth factor-reduced Matrigel (Corning) in conditioned media collected from control and miR-378aKO hiPSC-CM. Images of forming tubes were acquired using a Nikon Eclipse TS100 microscope four, six and 24 hours after seeding. Analysis of the total length of the tubes, number of nodes, number of branches and number of junctions was performed using Angiogenesis Analyzer for ImageJ software[10].

#### 1.5. Angiogenic potential of hiPSC-CM in organ-on-chip model

Analysis of the pro-angiogenic activity of control and miR-378aKO hiPSC-CM was performed using idenTx 3 microfluidic chips (AIM Biotech) as previously described by us[11] with modifications. Briefly, the gel channel of the chip was filled with 10 µL of hydrogel prepared by mixing bovine fibrinogen (Sigma-Aldrich) with thrombin from bovine plasma (Sigma-Aldrich). One of the lateral medium channels was filled with HAEC and the second one with either control or miR-378aKO hiPSC-CM in their respective media. Images of cell behavior in the following days of co-culture were acquired using a Nikon Eclipse TS100 microscope.

#### 1.6. Histological procedures

Upon perfusion hearts were mounted in OCT Compound (Leica) and frozen in liquid nitrogen–cooled isopentane or fixed for 48 h in 10% formalin and paraffin-embedded. Frozen (stored at –80 °C, cryostat sectioned) and/or paraffin (microtome sectioned) tissue samples (8 μm thick) were used for assessment of inflammatory cell infiltration and necrosis (H&E), fibrosis (Masson's trichrome staining, Sirius red staining) and immunofluorescent analysis. Histological procedures were done according to the vendor's instructions (Sigma–Aldrich). Immunofluorescent (IF) analysis of blood vessels (α-SMA+/CD31+/lectin binding) was done using previously established protocol for IF tissue stainings[12] with modifications described by us recently[13]. Slides were incubated with primary antibodies: rabbit anti-mouse α-smooth muscle actin (α-SMA) (1:200, Abcam, cat. no. ab5694), rat anti-mouse CD31 (1:150, BD Biosciences, cat. no. 550274), and lectin GSL-1 B4 DyLight 649 (15 μg/mL, Vector Labs, cat. no. DL-1208) in blocking buffer (overnight at 4 °C). This was followed by incubation with secondary antibodies: donkey anti-rabbit IgG Alexa Fluor 488 (Invitrogen, cat. no. A21206) and goat anti-rat IgG Alexa Fluor 568 (Invitrogen, cat. no. A11077), supplemented with 0.2 μg/mL DAPI. Images were taken with Leica DMi8 microscope with CMOS Leica MC170 HD camera and analyzed using Leica Application Suite X (Las X) Life Science microscope software platform as described previously[13].

#### 1.7. Periodic acid-Schiff (PAS) staining

To evaluate the level of glycogen content in control and miR-378aKO hiPSC-CM, PAS staining, adapted to cell culture from tissue staining[14] was performed. Briefly, cells were fixed for 10 min in 4% PFA, then stained for 10 min in 0.5% periodic acid-Schiff base aqueous solution and 20 min in periodic acid-Schiff base solution. After washing once, cells were counterstained for an additional 2 min with hematoxylin. Images were taken using a Nikon Eclipse TS100 microscope.

#### 1.8. Transmission electron microscopy (TEM) imaging

TEM imaging was performed as previously described[5]. Briefly, pellet of 100,000 cells was fixed with 2.5% glutaraldehyde in 0.1 M cacodylic buffer overnight at 4 °C, followed by 1% osmium tetroxide for 1 h at 4°C. Then, samples were dehydrated in graded ethanol 50%, 70%,

96% and 100%. After incubation in propylene oxide samples were embedded in Poly/Bed® 812 epoxy resin at 68 °C. In the next step, ultrathin sections, about ~70 nm thickness were cut using microtome, placed on 300-mesh Formvar/Carbon grids and contrasted using uranyl acetate and lead citrate. Imaging was done with a JEOL JEM 2100HT (Jeol Ltd, Tokyo, Japan) transmission electron microscope (TEM) that was used at an accelerating voltage of 80 kV. Images were taken by using 4 k × 4 k camera (TVIPS) equipped with EMMENU software ver. 4.0.9.87.

#### 1.9. RNA isolation, reverse transcription and quantitative PCR (qPCR)

Dissected murine hearts were preserved in RNA*later* tissue storage reagent (Sigma-Aldrich), snap-frozen in liquid nitrogen, and stored at –80 °C. For total RNA isolation tissues were homogenized using TissueLyser (Qiagen) in 1 mL of QIAzol Lysis Reagent (Qiagen) followed by organic extraction with 200 µL chloroform, and isopropanol precipitation. RNA quality and concentration was determined by NanoDrop Spectrophotometer (ThermoFisher Scientific).

Total RNA isolation from the cells was performed based on the Chomczyński method with modifications as previously described[1]. Cells were collected and washed 3 times with PBS, then covered with Fenozol reagent (400 µL, A&A Biotechnology) and chloroform (100 µL, POCH) was added. Samples were mixed vigorously for 1 minute, then incubated on ice for 20 min to separate the organic phase from the aqueous phase and centrifuged (10,000 × g, 4°C). The upper aqueous phase was collected in new tubes and an equal volume of cold isopropanol (POCH) was added and RNA was precipitated overnight at -20°C. The samples were then centrifuged (30 min, 10,000 × g, 4°C). The supernatant was discarded and the RNA pellet was washed twice with 70% ethanol (POCH). Subsequently, RNA was dried and dissolved in sterile water (15 µL). Secondary structures were destabilized by 10 min incubation at 65°C, then the concentration was measured using a NanoDrop spectrophotometer (ThermoFisher Scientific).

Reverse transcription (RT) of mRNA/miRNA was performed using 500 ng/10 ng of total RNA in a mix containing RevertAid reverse transcriptase (200 U/µL, ThermoFisher Scientific), dNTPs and oligodT (Genomed) or conducted using the miRCURY LNA RT kit (QIAGEN), respectively. RT was carried out in ProFlex PCR System (Applied Biosystems).

Gene expression analysis and miRNA detection were performed with SYBR Green JumpStart Taq Ready Mix (Sigma-Aldrich) or miRCURY LNA SYBR green PCR kit (Qiagen), respectively, and specific primers (Table 2). Quantitative PCR (qPCR) reaction was carried out in StepOne Plus Real-Time PCR System followed by data analysis using StepOne Software v2.3. A quantification of gene/miRNA expression in relation to *Eef2*/U6 snRNA reference was based on the comparative Ct method according to the $2^{-\Delta Ct}$ formula.

*Table 2. Sequence of primers used in the study (F – forward, R – reverse).*

| Gene | Primer sequence |
|---|---|
| *mhEEF2* | F: 5’ TCAGCACACTGGATAGAGG 3’ |
| | R: 5’ GACATCACCAAGGGTGTGCA 3’ |
| *TNNT2* | F: 5’ ATCCAGAACGCCCAGACAGA 3’ |
| | R: 5’ GCTGCTTGAACTTCTCCTGC 3’ |
| *NRF1* | F: 5’ GTGCTGATGAAGACTCGCCT 3’ |
| | R: 5’ GGCCGTTTCCGTTTCTTTCC 3’ |
| *SIRT1* | F: 5’ GTCAAGGGATGGTATTTATGCTCG 3’ |
| | R: 5’ GGCTATGAATTTGTGACAGAGAG 3’ |
| *TFAM* | F: 5’ CGAGGTGGTTTTCATCTGTC 3’ |
| | R: 5’ CTCTTCTTTATATACCTGCCACTC 3’ |
| *Col1a1* | F: 5’ CGATCCAGTACTCTCCGCTCTTCC 5’ |
| | R: 5’ ACTACCGGGCCGATGATGCTAACG 3’ |
| *Fn1* | F: 5’ AGCCTGCTCATCAGTTGGGA 3’ |
| | R: 5’ GATGGAAACTGGCTTGCTGC 3’ |
| *Vegfa* | F: 5’ ATGCGGATCAAACCTCACCAA 3’ |
| | R: 5’ TTAACTCAAGCTGCCTCGCCT 3’ |
| *Fgf1* | F: 5’ ATGGACACCGAAGGGCTTTT 3’ |
| | R: 5’ GAGGCCCACAAACCAGTTCT 3’ |
| *Kdr* | F: 5’ CGGCCAAGTGATTGAGGCAG 3’ |
| | R: 5’ ATGAGGGCTCGATGCTCGCT 3’ |
| *Cxcl12* | F: 5’ CCTTCAGATTGTTGCACGGCT 3’ |
| | R: 5’ CCCACCACTGCCCTTGCATC 3’ |
| *Cxcr4* | F: 5’ AAACCTCTGAGGCGTTTGGT 3’ |
| | R: 5’ AGCAGGGTTCCTTGTTGGAG 3’ |
| Sequences of primers used for the determination of miRNA level using miRCURY LNA PCR kit | |
| U6 snRNA | 5′ CGCAAGGATGACACGCAAATTC 3′ |
| miR-378a-3p | 5′ ACUGGACUUGGAGUCAGAAGG 3′ |
| hsa-miR-378a-5p | 5′ CUCCUGACUCCAGGUCCUGUGU 3′ |

#### 1.10. Protein isolation

Dissected cardiac tissues were snap-frozen in liquid nitrogen and stored at –80 °C until use. For protein extraction, tissues were homogenized at room temperature (rt) using a TissueLyser (Qiagen) in 500 µL of either 1% SDS in 0.1 M Tris-HCl (pH 7.5) for proteomic analysis, or in Pierce RIPA Buffer (Thermo Scientific) supplemented with a protease inhibitor cocktail. Homogenates were centrifuged at 20,000 × g (10 min, rt) for SDS buffer, or 8,000 × g (10 min, 4 °C) for RIPA buffer. The total protein concentration in 25-fold diluted supernatants was determined using either the Detergent Compatible Bradford Assay (Pierce) or the bicinchoninic acid (BCA) assay (Sigma-Aldrich), depending on buffer compatibility.

Total protein was isolated from hiPSC-CM by 30 min incubation of the washed cell pellet in RIPA buffer (Pierce) supplemented with protease inhibitors (Merck). Lysates were cleared of cell debris by 5 min centrifugation at 10,000 × g at 4 °C. Concentration was measured using BCA assay.

#### 1.11. Transcriptomic analysis

Transcriptomic analysis was performed as previously described[5]. Briefly, AmpliSeq™ Transcriptome Human Gene Expression Kit (Thermo Fisher Scientific) was used for the generation of RNA libraries, which were combined in equimolar amounts and sequenced on the Ion Proton™ Sequencer (Thermo Fisher Scientific) using the Ion PI™ Hi-Q™ Sequencing 200 Kit (ThermoFisher Scientific) and the Ion PI™ Chip Kit v3 (ThermoFisher Scientific). RNA samples collected from control hiPSC1-CM generated in five independent differentiations as well as RNA samples collected from hiPSC1-CM generated from three distinct miR-378aKO hiPSC1 clones (clone 16 – one differentiation, clone 44 and 48 – two independent differentiations) were used in this analysis. Bioinformatics analyses, including normalization, differential expression, and functional analysis were performed using DESeq2 (v1.40.1) and clusterProfiler (v4.8.3) packages[15,16] in R (v4.3.0).

#### 1.12. Proteomic analysis

Proteomic analysis of hiPSC-CM was performed as previously described[5]. Total protein extracts were prepared using 10% SDS (BioShop) in PBS. Samples were prepared by taking 10 µg of protein and diluted with PBS so that the final SDS concentration was 4%. Reduction and alkalization were performed by adding 5 mM Bond-Breaker TCEP (ThermoFisher Scientific) and 15 mM CAA (Chloroacetamide) (Sigma Aldrich) and heating for 10 min at 70°C. The prepared SP3 magnetic bead mixture (20 µL Sera-Mag A, 20 µl Sera-Mag B [SpeedBead Magnetic Carboxylate, Cytiva], 160 µL water) was placed on a magnet for 3 min, then the supernatant was collected, and the beads were washed twice with water and suspended in water. They were then added in a volume of 2 µL to the protein samples and acetonitrile was immediately added to a total concentration of 50%. After 8 min incubation at room temperature, the samples were placed on a magnetic stand for 2 min, then washed twice with 70% ethanol, once with acetonitrile and left to dry for 2 min. Samples were subjected to overnight (37 °C, 750 rpm) digestion with trypsin (0.1 µg, Serva) and LysC (0.1 µg, Wako). The next day, the samples were placed on a magnetic stand and the magnetic beads with bound proteins were suspended in buffer A (0.1% aqueous solution of formic acid) and the preparation of columns with a protein binding substrate (StageTip, double layer of SDB-RPS placed in a 0.2 mL pipette tip) was started by successive repetitions of centrifugation (1 min, 500 × g) and washing: with methanol, buffer B (elution buffer, 80% acetonitrile, 0.1% formic acid) and twice with buffer A. After transferring the samples to StageTip, the cycle of centrifugation (1 min, 500 × g) and washing was repeated, successively: twice with buffer B and buffer A. The measurements were eventually performed using a mass spectrometer (Multiscan FC, ThermoFisher). Proteomic analysis of murine hearts was performed using 5 µg/µl of cardiac protein lysates in 1% SDS 0.1 M Tris HCl pH 7.5 in Proteomics and Mass Spectrometry Core Facility of Malopolska Centre of Biotechnology, Krakow, Poland.

Preliminary proteomic data analysis was performed using MaxQuant[17] and Perseus[18] (1.6.15.0) software. It was followed by the differential expression analysis of $log_2$-transformed intensities with ANOVA in R (v4.3.0) and subsequent adjustment of p-values for multiple testing with Benjamini-Hochberg method. For hiPSC-CM, the chosen model included the donor's effect. To avoid any artifacts, no imputation of proteomic data was performed. Integrated analysis of transcriptomic and proteomic data from hiPSC-CM as well as proteomic data from murine hearts was performed using biomaRt[19] (v2.56.1).

#### 1.13. Western blot

Cardiac and hiPSC-CM protein samples (10-30 µg) were subjected to SDS–PAGE and Western Blot as described previously[1]. After 1 hour blocking in 5% BSA (bovine serum albumin) or 5% non-fat milk in 0.1% Tween 20 in TBS, membranes were incubated with primary antibodies (overnight at 4 °C), listed in Table 3. Secondary antibodies conjugated with HRP were added for 1 hour at room temperature [anti-rabbit (1:5000, Cell Signaling, cat. no. 7074P2) and anti-mouse (1:10,000, BD Pharmingen, cat. no. 554002)]. All antibodies were diluted in blocking buffer.

*Table 3. Primary antibodies used in the study.*

| Antibody | Dilution | Company | Catalog number |
|---|---|---|---|
| AKT | 1:500, 5% BSA | Cell Signaling | #9272 |
| pAKT | | | #9271S |
| PGC1b | | Abcam | ab176328 |
| PGC1α | | | ab54481 |
| IGF1R | | Santa Cruz Biotechnology | sc713 |
| SIRT1 | | Abcam | ab110304 |
| GLUT1 | 1:20 000, 5% milk | | ab115730 |
| NRF1 | 1:500, 5% milk | Cell Signaling | #46743S |
| pERK1/2 | 1:1000, 5% milk | | #5301 |
| ERK1/2 | 1:500, 5% milk | | #9102 |
| PathScan Multiplex Western Cocktail I | 1:1000, 5% BSA | | #5301 |
| TFAM | 1:500, 5% BSA | | #7495S |
| pAMPKα | | | #2535 |
| AMPKα | | | #5831T |
| AMPKγ | | | #4187 |
| Optineurin (OPTN) | | | #58981 |
| GSK3α/β | | Santa Cruz Biotechnology | sc-9166 |
| pGSK3α/β | | | sc-135653 |
| p62 | 1:500, 5% milk | Cell Signaling | #8025 |
| Parkin | | | #4211 |
| LC3b I/II | | | #3868 |
| GAPDH | | Santa Cruz Biotechnology | sc-59540 |
| α Tubulin | | Sigma-Aldrich | T9026 |

#### 1.14. ELISA

Detection of murine FGF2 (R&D Systems, #DY3139) and VEGFA (R&D Systems, #DY3139) protein in heart lysates and human VEGF (R&D Systems, #DY293B-05) in hiPSC-CM-conditioned culture media was performed according to vendor's protocols.

#### 1.15. Immunofluorescent analysis

Immunocytochemical staining was performed on cells fixed with 4% paraformaldehyde (PFA), which were then washed three times with PBS, permeabilized in 0.1% TRITON X100 in PBS for 20 min and washed again. Blocking was performed by a 1 h incubation in 3% BSA. Primary antibodies (anti-OCT4, 1:200, sc-33759, Santa Cruz Biotechnology; anti-NANOG, 1:100, sc-293121, Santa Cruz Biotechnology; anti-SSEA4, 1:200, sc-21704, Santa Cruz Biotechnology; anti-TRA-1-60, 1:200, MAB4360, Millipore; anti-TRA-81, 1:200, MAB4381, Millipore; anti-TFAM, 1:200, 7495S, Cell Signaling) were diluted in 3% BSA and added to the cells for overnight incubation at 4°C. After washing, fluorochrome-conjugated secondary antibodies (Alexa Fluor 568 donkey anti-mouse, Alexa Fluor 488 donkey anti-goat, Alexa Fluor donkey anti rabbit 488, Invitrogen) were added at a 1:400 dilution in 3% BSA for one hour. Cell nuclei were stained with Hoechst 33342 (1 µg/mL). Images were taken with a Nikon Eclipse TS100 microscope or a Carl Zeiss LSM-510 meta laser scanning confocal microscope at 200x and 400x magnification, respectively.

#### 1.16. Mitochondrial content analysis

Mitochondrial content was determined by the ratio of mitochondrial DNA (mtDNA) to nuclear DNA, verified by qPCR performed as for gene expression analysis, using the following primer: gDNA forward - 5' CTATGGGACGCTTGATGT 3', gDNA reverse - 5' GCAATCATTCGTCTGTT 3', mtDNA forward - 5' CCCTAAAACCCGCCACATCT 3', mtDNA reverse - 5' GAGCGATGGTGAGAGAGCTAAGGT 3'. DNA was isolated using Genomic Mini kit (A&A Biotechnology) according to the manufacturer's protocol. 30 ng of isolated DNA was then used as a template in qPCR analysis.

#### 1.17. Glucose uptake

Glucose uptake was determined using the Glucose Uptake-Glo™ Assay (Promega). Briefly, 10,000 cells were plated in a 96-well plate dedicated for bioluminescence measurements. On the day of the assay, hiPSC-CM were washed with PBS and 50 µL of 1 mM 2DG (2 deoxyglucose) was added and incubated at RT for 10 min, followed by 25 µL of stop buffer and an equal volume of neutralization buffer. After adding 100 µL of detection buffer and 30 min incubation, the luminescence measurement was performed using an Infinite M200 (Tecan) plate reader.

#### 1.18. Hexokinase activity measurement

Hexokinase activity was measured colorimetrically using Hexokinase Activity Assay Kit (Abcam). Briefly, 10,000 cells were harvested, washed, and suspended in appropriate lysis buffers according to the manufacturer's instructions.

#### 1.19. Lactate dehydrogenase (LDH) activity measurement

LDH activity was measured colorimetrically using Lactate Dehydrogenase Assay Kit (Abcam). Briefly, 10,000 cells were harvested, washed, and suspended in appropriate lysis buffers according to the manufacturer's instructions.

***Supplementary Table 1.*** *The results of transcriptomic differential expression analyses of miR-378aKO hiPSC-CM in comparison to their control counterparts. The analysis was performed using DESeq2 (v1.40.1) package in R (v4.3.0). The NA values in the "padj" column result from the independent filtering performed by DESeq2 prior to the adjustment for multiple testing (see DESeq2 manual).*

***Supplementary Table 2****. The results of GO-terms over-representation analysis of miR-378aKO hiPSC-CM transcriptomic data in comparison to their control counterparts. The analysis was performed using clusterProfiler (v4.8.3) package in R (v4.3.0). Only records with adjusted p-value < 0.05 are shown.*

***Supplementary Table 3.*** *The results of KEGG pathways over-representation analysis of miR-378aKO hiPSC-CM transcriptomic data in comparison to their control counterparts. The analysis was performed using clusterProfiler (v4.8.3) package in R (v4.3.0). Only records with adjusted p-value < 0.05 are shown.*

***Supplementary Table 4.*** *The results of GO-terms over-representation analysis of combined transcriptomic and proteomic dataset from control and miR-378aKO hiPSC-CM as well as proteomic dataset from miR-378a+/+ and miR-378a-/- hearts of 11-12-week-old mice. Only genes with a consistent direction of expression change across all three datasets were included in the analysis. The analysis was performed using clusterProfiler (v4.8.3) package in R (v4.3.0). Only records with adjusted p-value < 0.05 are shown.*

***Supplementary Table 5.*** *The results of KEGG pathway over-representation analysis of combined transcriptomic and proteomic dataset from control and miR-378aKO hiPSC-CM as well as proteomic dataset from miR-378a+/+ and miR-378a-/- hearts of 11-12-week-old mice. Only genes with a consistent direction of expression change across all three datasets were included in the analysis. The analysis was performed using clusterProfiler (v4.8.3) package in R (v4.3.0). Only records with adjusted p-value < 0.05 are shown.*
